# *In vivo* tests of the *E. coli* TonB system working model—interaction of ExbB with unknown proteins, identification of TonB-ExbD transmembrane heterodimers and PMF-dependent ExbD structures

**DOI:** 10.1101/2024.07.10.602958

**Authors:** Kathleen Postle, Dale Kopp, Bimal Jana

## Abstract

The TonB system of *Escherichia coli* resolves the dilemma posed by its outer membrane that protects it from a variety of external threats, but also constitutes a diffusion barrier to nutrient uptake. Our working model involves interactions among a set of cytoplasmic membrane-bound proteins: tetrameric ExbB that serves as a scaffold for a dimeric TonB complex (ExbB_4_-TonB_2_), and also engages dimeric ExbD (ExbB_4_-ExbD_2_). Through a set of synchronized conformational changes and movements these complexes are proposed to cyclically transduce cytoplasmic membrane protonmotive force to energize active transport of nutrients through TonB-dependent transporters in the outer membrane (described in Gresock et *al.*, J. Bacteriol. 197:3433). In this work, we provide experimental validation of three important aspects of the model. The majority of ExbB is exposed to the cytoplasm, with an ∼90-residue cytoplasmic loop and an ∼50 residue carboxy terminal tail. Here we found for the first time, that the cytoplasmic regions of ExbB served as *in vivo* contacts for three heretofore undiscovered proteins, candidates to move ExbB complexes within the membrane. Support for the model also came from visualization of *in vivo* PMF-dependent conformational transitions in ExbD. Finally, we also show that TonB forms homodimers and heterodimers with ExbD through its transmembrane domain *in vivo*. This trio of *in vivo* observations suggest how and why solved *in vitro* structures of ExbB and ExbD differ significantly from the *in vivo* results and submit that future inclusion of the unknown ExbB-binding proteins may bring solved structures into congruence with proposed *in vivo* energy transduction cycle intermediates.

## INTRODUCTION

The outer membranes of Gram-negative bacteria provide protection from harmful agents such as detergents and antibiotics, but limit nutrient access to those that can be obtained by diffusion through β-barrel porins [size cut-off of ∼600 Da for *Escherichia coli*); (1)]. They circumvent the limitations imposed by the outer membrane using proteins integral to both the cytoplasmic and outer membranes, the aggregate being termed the TonB system.

The known *E coli* TonB system consists of TonB (239 residues), ExbB (244 residues), and ExbD (141 residues) located in the cytoplasmic membrane and various TonB-dependent transporters (TBDTs) in the outer membrane (Fig.1). For *E. coli*, the transport ligands are cobalamin from exogenous sources and siderophores synthesized and excreted by the *E. coli* itself, other bacteria, and fungi into the environment to capture iron (2). The four known components of the TonB system in Fig. 1 are all regulated by iron availability (3–6). The siderophore enterochelin (a.k.a enterobactin), the only siderophore that is synthesized by *E. coli* itself, is transported across the outer membrane by the TBDT FepA (7–9). Other Gram-negative bacteria use the TonB system for a variety of other nutrients that are too large, too scarce, or too important to rely on diffusion through the outer membrane porins (10). Because some are valuable to hosts, TonB systems are virulence factors for pathogenic Gram-negative bacteria (11). The relative simplicity of the *E. coli* TonB system, with a single set of *exbB/exbD* and *tonB* genes makes it a useful model for study. In comparison, there are 8 sets of predicted *tonB-exbB-exbD* genes in *Xanthomonas* and 120 predicted TBDTs in *Bacterioides* (10).

**Figure 1.**
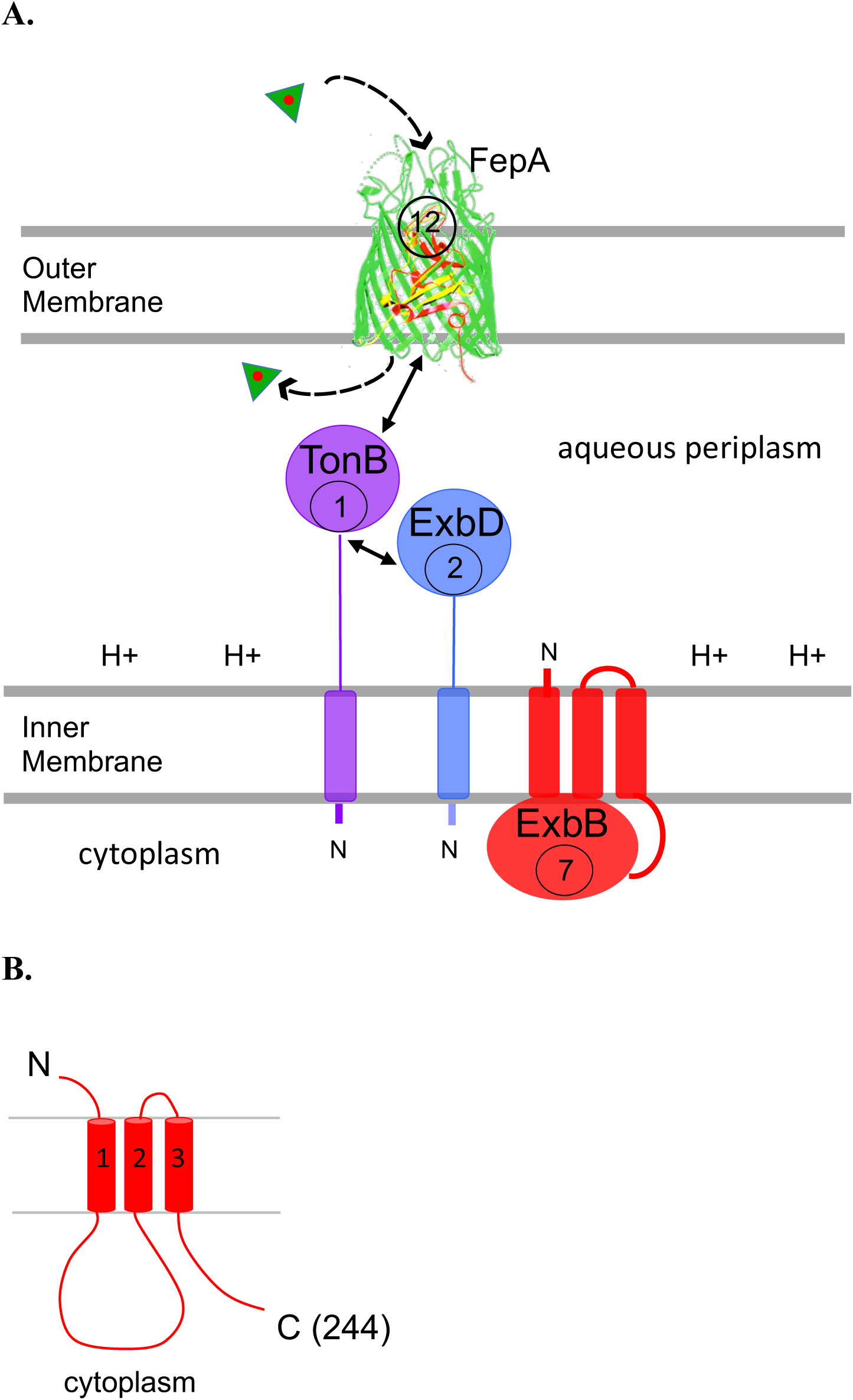
The TonB system of *Escherichia coli* K12. **A.)** The TonB-dependent transporter, FepA, is shown in the outer membrane. The cytoplasmic (inner) membrane protein ExbD uses the protonmotive force of the cytoplasmic membrane (H+) to configure the periplasmic domain of TonB, which binds to FepA and induces active transport of siderophore enterochelin (green triangle with red center) through FepA and into the periplasm. Ratios of the four proteins *in vivo* are indicated in circles in each protein. FepA is by far the most abundant, suggesting that there must be an energy transduction cycle where TonB binds and releases FepA (6). Our current model for the molecular details of the energy transduction cycle is shown in Fig. 2. The crystal structure of FepA was solved by Buchanan et al. (96). **B.)** ExbB has type III topology: N out, C in (97). The ExbB cytoplasmic loop domain between transmembrane domains 1 and 2 is predicted to contain residues ∼43-131. The cytoplasmic carboxy terminal tail following transmembrane domain 3 is predicted to contain residues ∼197-244 (31).

The molecular mechanism by which TonB/ExbB/ExbD participate in transducing cytoplasmic membrane proton motive force (PMF) energy to a TBDT for active transport of extra-cellular ligands across the outer membrane (12) remains a mystery. All models agree that TonB directly contacts the TBDTs to decrease their high affinity for ligands (sub-nanomolar in many cases), enabling their vectoral release into the periplasmic space. However, initial high affinity binding of ligands to TBDTs is independent of both PMF and the TonB system, while subsequent active transport across the outer membrane via the TBDT requires each (13–15). Adding to the mystery, the periplasmic domain of TonB binds TBDTs *in vivo* and *in vitro* whether or not it is “energized” by PMF or even active (16–21).

The number of TonB molecules per cell is much lower than the number of TBDTs, indicating the need for an energy transduction cycle where TonB cyclically binds and releases TBDTs for productive transport of ligands into the periplasm (6). Consistent with that, ligand-loaded TBDTs compete for TonB *in vivo* (22). Once in the periplasm, siderophores such as enterochelin are scavenged by a periplasmic binding protein and delivered to an ABC transporter for TonB-<u>in</u>dependent transport into the cytoplasm (23) (not shown in Fig. 1).

ExbB serves as a scaffold for both TonB and ExbD and is required for their proteolytic stability. Consistent with that role, ExbB itself is proteolytically stable, even when overexpressed (5, 19, 24–28). It is a member of the *E. coli* ExbB-TolQ-MotA family of cytoplasmic membrane proteins, which have similar topologies, residue similarities in their distal two transmembrane domains, and interact with their respective cytoplasmic membrane proteins, ExbD-TolR-MotB, each containing an essential Asp residue within their transmembrane domains (29–31). Using the PMF, TolQ/R play a role in cell division and MotA/B play a role in energizing the flagellar motor (32, 33).

While the role that ExbB plays in the TonB system is becoming clearer, we find no evidence that its three transmembrane domains directly participate in proton translocation *in vivo* (31). Rather, ExbB transmembrane domain 1, the least conserved of the three transmembrane domains, appears to be the primary contact with the TonB transmembrane domain, consistent with the finding that three different ExbB suppressors of TonB transmembrane domain mutations have been isolated in ExbB transmembrane domain 1 [Fig. 1B); (31, 34, 35)]. The suppressors are not allele-specific and substitute Asp or Glu residues for aliphatic residues. Similar TolQ suppressors exist in the Tol system (36, 37). ExbB and TonB also formaldehyde-cross-link *in vivo* (19, 31).

Specific residues in the ExbB cytoplasmic carboxy terminus distal to transmembrane domain 3 are required for PMF-dependent interaction between TonB and ExbD periplasmic domains, suggesting that signals are being transduced between the cytoplasm and the periplasm (19, 38). ExbB transmembrane domains 2 and 3, which boundary its short periplasmic loop, appear to play roles in mediating that signal transduction (31).

Results suggest that a full complement of proteins in the TonB system has yet to be defined. Cytoplasmic domains of MotA and non-*E. coli* members of the ExbB-TolQ-MotA family bind cytoplasmic proteins that modulate their activities, suggesting that ExbB might do likewise (39–41). ExbB formaldehyde cross-links *in vivo* as a dimer of dimers (tetramer) bound to ∼85 kDa of unknown protein, a complex called tetramer + X (38). In addition, cessation of protein synthesis leads to rapid loss of TonB system activity (T_1/2_ ≅ 20 min), while each of the four known proteins, TonB, ExbB, ExbD and FepA has a chemical half-life of ≅ 120 min, suggesting the existence of at least one unknown protein in the TonB system with a short chemical half-life (27, 42).

Like ExbB transmembrane domains, the role of the TonB transmembrane domain in PMF response has also been ruled out (43). Thus, ExbD and its transmembrane domain residue Asp 25 are the sole PMF-responsive components in the TonB system (19, 44). The negative charge and position of Asp 25 are virtually essential. Conservative replacement with Glu at ExbD residue 25 decreases TonB system activity to 5-10% of the wild-type value, whilst replacements with all other residues inactivates ExbD. Likewise, the Asp 25 residue cannot be relocated to another position in the transmembrane domain and support activity (45). ExbD appears to use cytoplasmic membrane PMF to correctly configure the TonB periplasmic carboxy terminus for transmission of energy to FepA (19, 46, 47).

Both ExbD and TonB can be functionally divided into two domains. Their uncleaved amino termini anchor them in the cytoplasmic membrane while most of their residues occupy the periplasmic space (Fig. 1). The TonB amino terminus contains a signal sequence used for export and required for its activity, even when the carboxy terminus is exported to the periplasm by other means (48). The TonB carboxy terminus undergoes dynamic conformational changes during an energy transduction cycle, interacting with another TonB carboxy terminus, with the carboxy terminus of ExbD, and with TBDTs or their phage/colicin infectious agents using nearly the same set of residues (18, 42, 49–54). Phenotypes and disulfide formation by Cys substitutions in the homodimerized TonB carboxy terminus show that solved structures of TonB homodimers (55, 56) do not exist *in vivo* (50). The periplasmic domain of ExbD is also conformationally dynamic, forming both TonB heterodimers and ExbD homodimers (51).

We had previously proposed a working model for the TonB system consistent with data from *in vivo* experiments [Fig. 2; (42)]. The present study validates and extends important aspects of the model. 1) First, the model predicts the existence of previously unknown cytoplasmic proteins that interact with ExbB, a key player in the TonB system. We show for the first time that an unknown ∼34 kDa protein binds to the ExbB cytoplasmic loop domain, and two additional unknown proteins of ∼24 kDa and ∼29 kDa bind to the cytoplasmic carboxy terminus *in vivo*. 2) Further, it predicts that TonB homodimerizes through its transmembrane domain and also heterodimerizes with the ExbD transmembrane domain during an energy transduction cycle, hypotheses that we test and validate here. 3) In addition, it predicts that the structure of the periplasmic domain of ExbD is dependent upon its ability to respond to PMF, an additional hypothesis that we test and validate here. Taken together, the results suggest both how and why there are major differences between models reflecting *in vivo* results and the solved structures of ExbB/D. A potential resolution of the differences is suggested.

**Figure 2.**
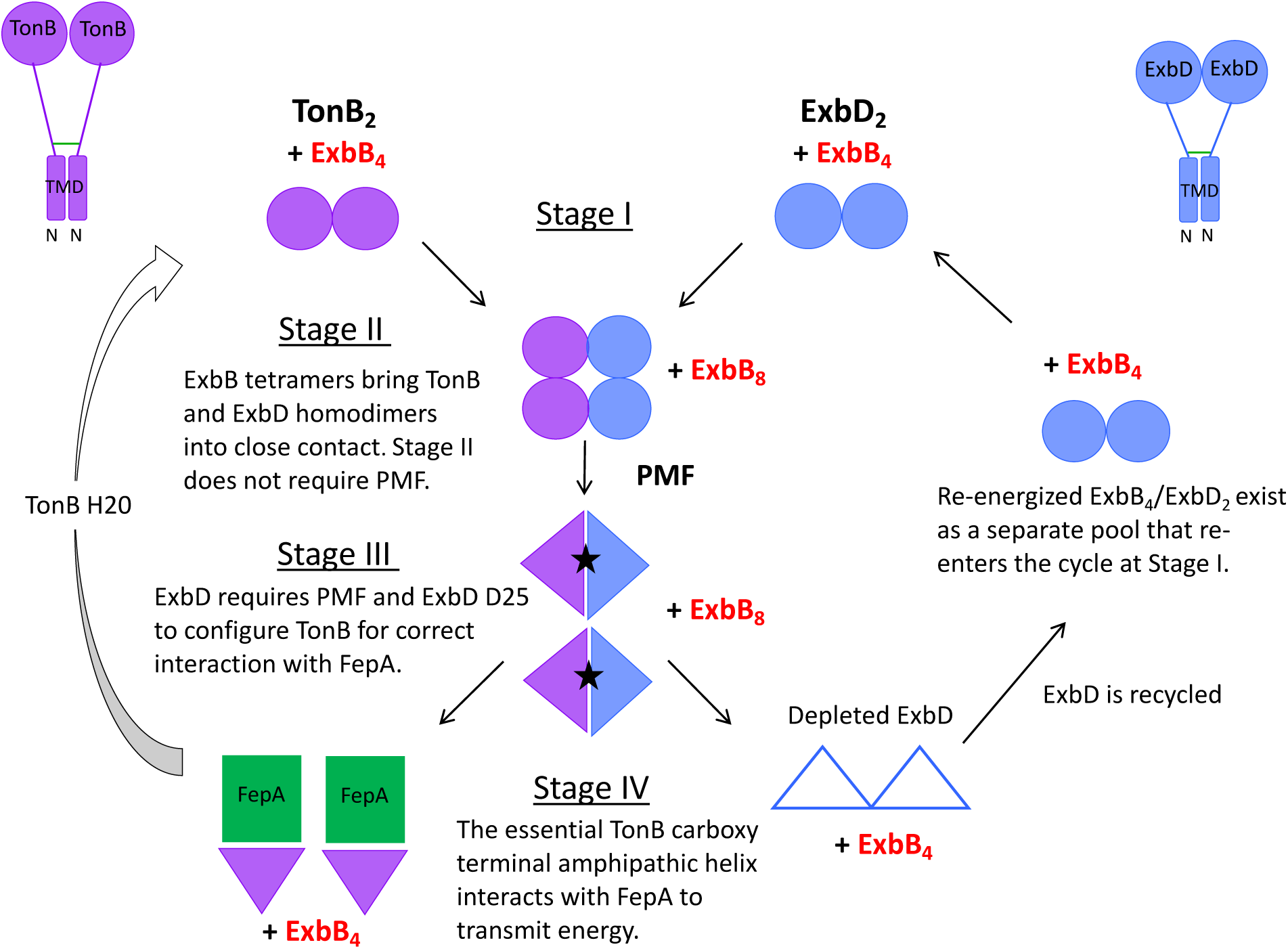
Model for the dynamic role of ExbB in the TonB energy transduction cycle based on *in vivo* data [adapted from (42)]. The model requires that one or more unknown cytoplasmic proteins transit ExbB tetramers associated with TonB and ExbD through the different stages. Full length TonB (purple, upper left corner) and full-length ExbD (blue, upper right corner) each have a single transmembrane domain signal anchor in the cytoplasmic membrane with the bulk of the residues occupying the periplasmic space (Fig. 1). TonB homodimerized near its transmembrane domain (green line) remains active throughout the cycle, suggesting it functions as a homodimer during energy transduction (42). The ExbD transmembrane domain also homodimerizes [green line; (45)], but it is not known if the homodimer persists throughout the cycle. In this model, only interactions between the periplasmic carboxy-terminal domains of TonB_2_ and ExbD_2_ homodimers are shown. Purple circles/triangles represent TonB carboxy termini. All stages of the TonB energy transduction cycle involve the participation of the essential carboxy terminal amphipathic helix (residues 199-216) in the TonB carboxy terminus (18, 52, 54). Blue circles/triangles represent ExbD carboxy termini. ExbB tetramers identified *in vivo,* ExbB_4_ in this model, independently stabilize both TonB and ExbD (24, 38). In **Stage I**, the carboxy terminal TonB periplasmic domains form obligatory homodimers through residues in and near the amphipathic helix (residues 199-216) [left of center; (18, 42, 50)]. ExbD periplasmic domains also homodimerize [right of center; (51)]. There is no evidence that TonB and ExbD homodimers interact at this stage. Because both are proteolytically stabilized by ExbB, it suggests that they exist as homodimers with separate ExbB tetramers. In **Stage II** (beneath Stage I), TonB and ExbD periplasmic domain homodimers are brought into close contact by movement of their respective ExbB tetramers such that ExbD now protects TonB from degradation by exogenously added protease in spheroplasts (35, 47, 51). In **Stage III** (beneath Stage II), PMF mediated through ExbD D25 configures the ExbD periplasmic domain, which configures the TonB periplasmic domain such that it can transmit energy to FepA (19, 46, 47, 51, 52). To accomplish this, the previously homodimeric TonB and ExbD periplasmic domains are converted into TonB-ExbD heterodimers, resulting in a TonB conformation that is now sensitive to degradation by exogenously added protease in spheroplasts (47, 51). Neither the TonB nor ExbB transmembrane domains appear to participate in the response to PMF (31, 43). **Stage IV** (left), the amphipathic helix of TonB binds to the TonB-dependent transporter, FepA (represented by the green square) (54). The siderophore enterochelin is actively transported across the outer membrane into the periplasmic space. The TonB transmembrane domain residue H20 provides a signal required for re-establishment of the obligatory ExbB_4_-TonB_2_ carboxy terminal homodimers in Stage I (42, 43, 50). **Stage IV** (right), after a transport event, energized ExbB_4_-ExbD_2_ is depleted (open triangles) and subsequently replenished by an unknown mechanism. A cellular ratio of 14 ExbB: 4ExbD: 2TonB [reflecting that TonB can remain homodimerized near its amino terminus (42) and through its amino terminus (this study)] is found in bacteria grown under three different conditions of iron availability (6). Similar ratios have been observed for the Tol system (82, 83). A separate pool of replenished ExbB_4_-ExbD_2_ is hypothesized to exist to bring the ratio in the model to 12ExbB:4ExbD:2TonB. We speculate that the separate pool is to provide “energized” ExbB_4_-ExbD_2_ homodimers that initiate the energy transduction cycle in Stage I. To account for the full cellular ratio, there may also be a small pool of free ExbB_4_. See (42) for a more complete explanation of the experimental basis for the model.

## RESULTS

### The ExbB cytoplasmic loop

#### All 10-Ala substitutions eliminate immediate growth arrest, and some exhibit modest activity

Because most of its residues occupy the cytoplasm, ExbB is the primary means by which the TonB system can respond to cytoplasmic signals, while only the amino terminal ∼12 and ∼22 residues of TonB and ExbD respectively access the cytoplasm (Fig. 1A). The ExbB cytoplasmic loop domain between transmembrane domains 1 and 2 (residues ∼43-131) and cytoplasmic carboxy terminus following transmembrane domain 3 (residues ∼197-244) are essential for its activity [Fig. 1B; (28, 31, 38)].

Eight of nine plasmid-encoded 10-residue deletions (Δ10s) of the ExbB cytoplasmic loop cause immediate, dominant, but reversible, growth arrest when induced with sufficient arabinose to reach chromosomal levels [Table 1, showing data from (28)]. The level of arabinose required provides a rough measure of sensitivity to endogenous proteases, since ExbB itself is a stable protein and requires little added arabinose to achieve the steady state chromosomal levels of expression encoded by strain W3110 (24, 28, 38). Immediate growth arrest is not due to collapse of the PMF and does not require the presence of ExbD or TonB, indicating that it is intrinsic to each ExbB variant and any other proteins with which it might interact (28). Taken together, the data suggest that growth arrest may reflect a changed interaction between the ExbB cytoplasmic loop and one or more unknown cytoplasmic growth-regulatory proteins (28).

**Table 1.** Comparison of Δ10 and 10-Ala variants in the ExbB cytoplasmic loop.

**Comparison of $\Delta 10$ and 10-Ala variants in the ExbB cytoplasmic loop**
|  | % Arabinose for<br>Chromosomal Expression<sup>a</sup> |  | Activity<sup>b</sup> |  | Immediate Growth<br>Arrest<sup>c</sup> |  |
| --- | --- | --- | --- | --- | --- | --- |
| | $\Delta 10^d$ | 10-Ala | $\Delta 10^d$ | 10-Ala | $\Delta 10^d$ | 10-Ala |
| Wild-type<sup>e</sup> | 0.00025% | 0.0005% | 100% | 100% | no | no |
| 40-49<sup>f</sup> | 0.1% | 0.002% | 0% | 0% | yes | no |
| 50-59 | 0.1% | 0.2% | 0% | 0% | yes | no |
| 60-69 | 0.03% | 0.2% | 0% | 0% | yes | no |
| 70-79 | 0.001% | 0.002% | 0% | ~30% | yes | no |
| 80-89 | 0.1% | 0.2% | 0% | 0% | yes | no |
| 90-99 | 0.001% | 0.002% | 0% | 0% | yes | no |
| 100-109 | 0.00025% | 0.003% | ~25% | ~10% | no | no |
| 110-119 | 0.1% | 0.2% | 0% | 0% | yes | no |
| 120-129 | 0.1% | 0.001% | 0% | 100% | yes | no |

<sup>a</sup>An indirect measure of sensitivity to endogenous proteases. The higher the level of arabinose required to achieve chromosomally encoded levels of expression, the greater the proteolytic sensitivity.
<sup>b</sup>Activity measured as a % of initial rates of iron transport compared to wild-type W3110. Effects of the $\Delta 10$ and 10-Ala substitutions were measured in the RA1017 strain background (W3110 $\Delta exbBD::kan$ , $\Delta tolQRA$ ).
<sup>c</sup>Growth arrests immediately upon addition of 0.1% or 0.2% arabinose as seen in (28). See Fig. 3 for 10-Ala data.
<sup>d</sup>Data for cytoplasmic loop $\Delta 10$ s is from (28).
<sup>e</sup>Wild-type is defined as plasmid pKP660 in RA1017.
<sup>f</sup>Numbers refer to the *exbB* codons that were modified in plasmid pKP660.

To gauge the importance of the cytoplasmic loop domain in the absence of the large structural perturbations caused by deleting 10-residue blocks, we engineered substitutions of ten Alanine residues (10-Alas) at corresponding deletion sites in plasmid pKP660. Most of the 10-Alas were largely inactive (<10% of wild-type) in iron transport assays, suggesting that residues essential for activity had been replaced and confirming the importance of the ExbB cytoplasmic loop (Table 1). The three exceptions, judged to be somewhat proteolytically stable, were: 1). ExbB 70-79Ala which exhibited ∼30% of wild-type activity; 2). ExbB 100-109Ala, which exhibited ∼10% of wild-type activity; 3). ExbB 120-129Ala which retained 100% of wild-type activity even though it lacked the wild-type V_122_GRQMGRG_129_ sequence, suggesting that none of its corresponding wild-type residues were functionally important. ExbB 50-59Ala, 60-69Ala, 80-89Ala, and 110-119Ala all required the maximal amount of arabinose to reach chromosomal levels, indicating that they were proteolytically highly unstable. Importantly, despite being inactive and proteolytically unstable like their Δ10 counterparts, none of the 10-Ala substitutions exhibited immediate growth arrest (Table 1; Figs. 3, 4A).

**Figure 3.**
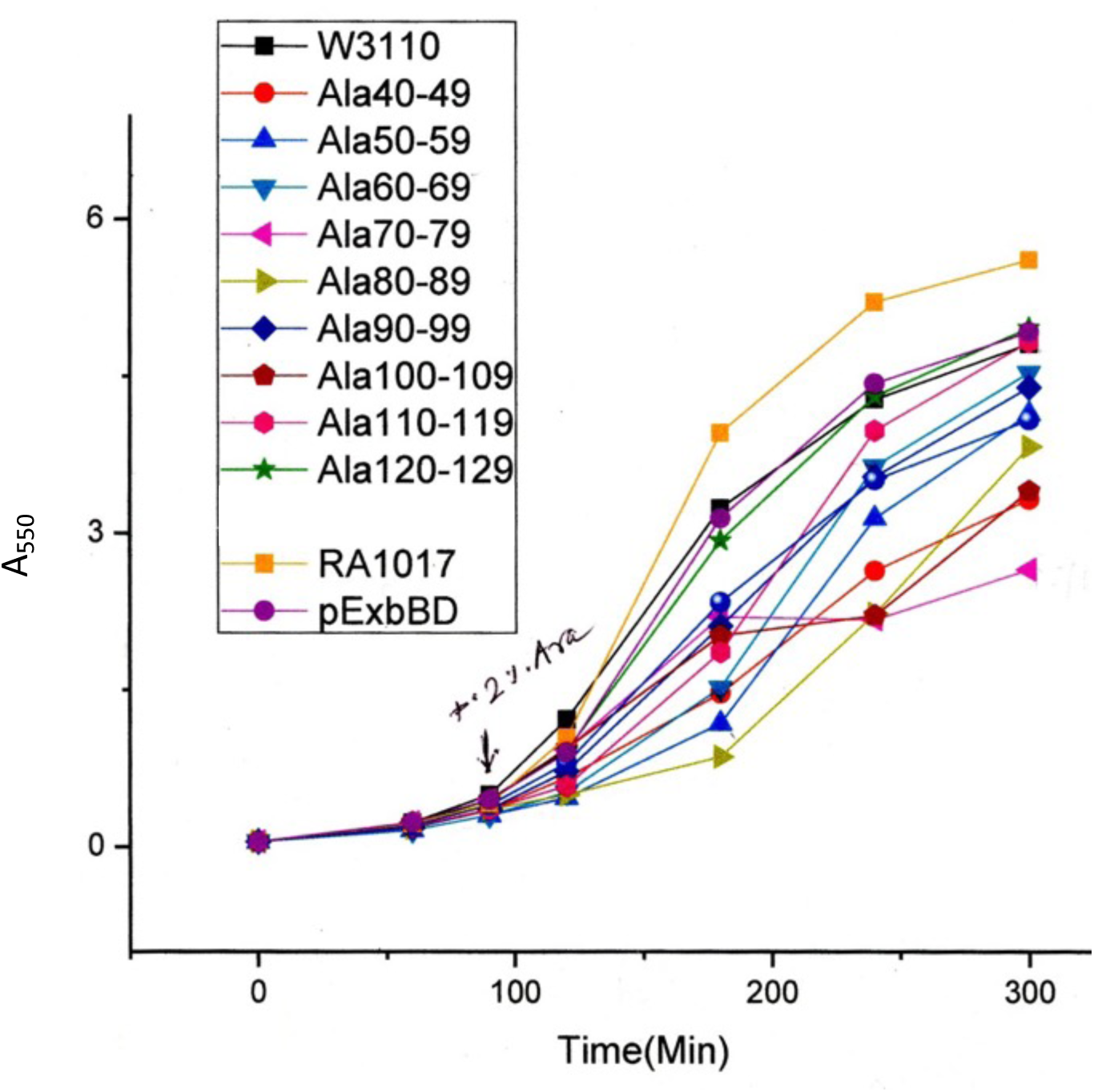
None of the 10Ala substitutions in the ExbB cytoplasmic loop exhibits immediate growth arrest. RA1017 (W3110 Δ*exbB/D*, Δ*tolQRA*) carrying various plasmids from Table 4 was grown overnight to saturation in LB plus 100 µg/ml ampicillin and then sub-cultured 1:100 into the same medium. After 1.5 hr, arabinose to 0.2% (maximum induction) was added and the absorbance of each culture at 550 nm was recorded for 3.5 hr. Variants examined are identified in the box. Immediate growth arrest was defined as lack of culture growth following 0.2% arabinose addition and for the remainder of the assay period (28). All plasmids are derivatives of pKP1657 (pExbB/D).

**Figure 4.**
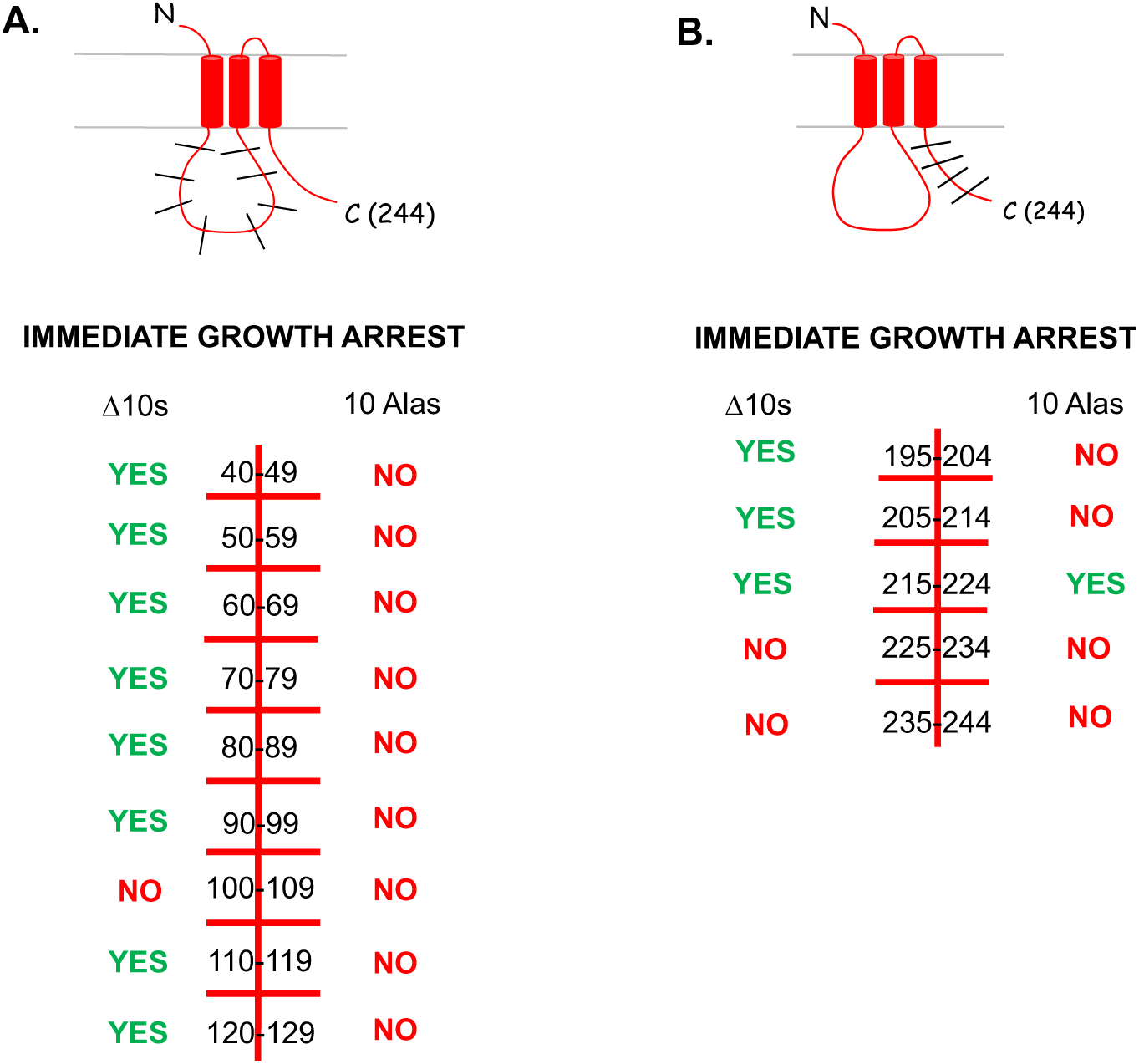
Summary of immediate growth arrest phenotypes for ExbB Δ10s and 10Alas. “Yes” indicates that immediate growth arrest occurred. “No” indicates that immediate growth arrest did not occur. **A. ExbB Cytoplasmic Loop**. Residues modified are listed in the center of the table with phenotypes of ExbB Δ10 variants on the left and ExbB 10-Ala substitutions on the right. Data for ExbB Δ10 variants come from (28). Data for the corresponding 10-Alas come from Fig. 3. **B. ExbB Cytoplasmic Carboxy Terminal Tail.** Residues modified are listed in the center of the table with phenotypes of ExbB Δ10 variants on the left and ExbB 10-Ala substitutions on the right. Growth arrest data are from Table 3. The topology and sections of ExbB described in the phenotype summaries are shown above each table.

#### ExbB forms a ∼60 kDa complex with an unknown protein through its cytoplasmic loop domain

A review on TonB concluded in 1990: “It is therefore premature to conclude that all of the auxiliary proteins that modulate TonB function have been genetically identified. TonB-dependent transport appears to be a finely balanced, multicomponent system. Until all of the ‘players’ have been identified and overexpressed, results from studies in which only a subset are overexpressed should be interpreted with caution” (75). Here we used *in vivo* photo-cross-linking to determine if unknown proteins bound to cytoplasmic loop residues of ExbB (57).

Unlike the thermolabile *in vivo* formaldehyde cross-linking (19, 38), *in vivo* photo-cross-linking results in protein-protein covalent bonds that are impervious to boiling in SDS polyacrylamide gel sample buffer (58). Both techniques probe for closely interacting proteins. For formaldehyde, reactive amino acids must be within 2.3–2.7 Å, and for photo-cross-linking, within 3 Å (59, 60). A second advantage of *in vivo* photo-cross-linking is that, unlike formaldehyde cross-linking, the residue through which a cloned protein initiates contact can be stipulated. We have highly specific antibodies against all three known cytoplasmic membrane proteins in the TonB system--ExbB, ExbD and TonB (6) --which allow identification of known partner proteins based on antibody reactivity, the apparent mass of the complex, and previous studies of formaldehyde cross-linked complexes (19, 38, 61). Photo-cross-linked ExbB complexes containing unknown proteins were identified as having anomalous apparent masses not previously observed and that did not react with either TonB or ExbD antibodies.

To search for unknown proteins that interacted with the ExbB cytoplasmic loop, we replaced *exbB* F48, F78, F113, and Q125 codons in plasmid-encoded *exbB/D* (pKP1657) with amber codons and expressed them in the presence of plasmid pEVOL-Bpa and the photo-cross-linkable amino acid pBpa (*p*-benzoyl-l-phenylalanine) in RA1017 (W3110, Δ*exbBD*::*kan*, Δ*tolQRA*). Following illumination with UV light, we detected an ExbB-specific, UV-specific complex of ∼60 kDa for F78 (pBpa), F113 (pBpa), and Q125 (pBpa) (Fig. 5A, lanes 4, 6, and 8) in addition to full-length monomeric ExbB. Given the calculated mass of ExbB (26.3 kDa), the results suggested that an unknown protein of ∼34 kDa was in the complex.

**Figure 5.**
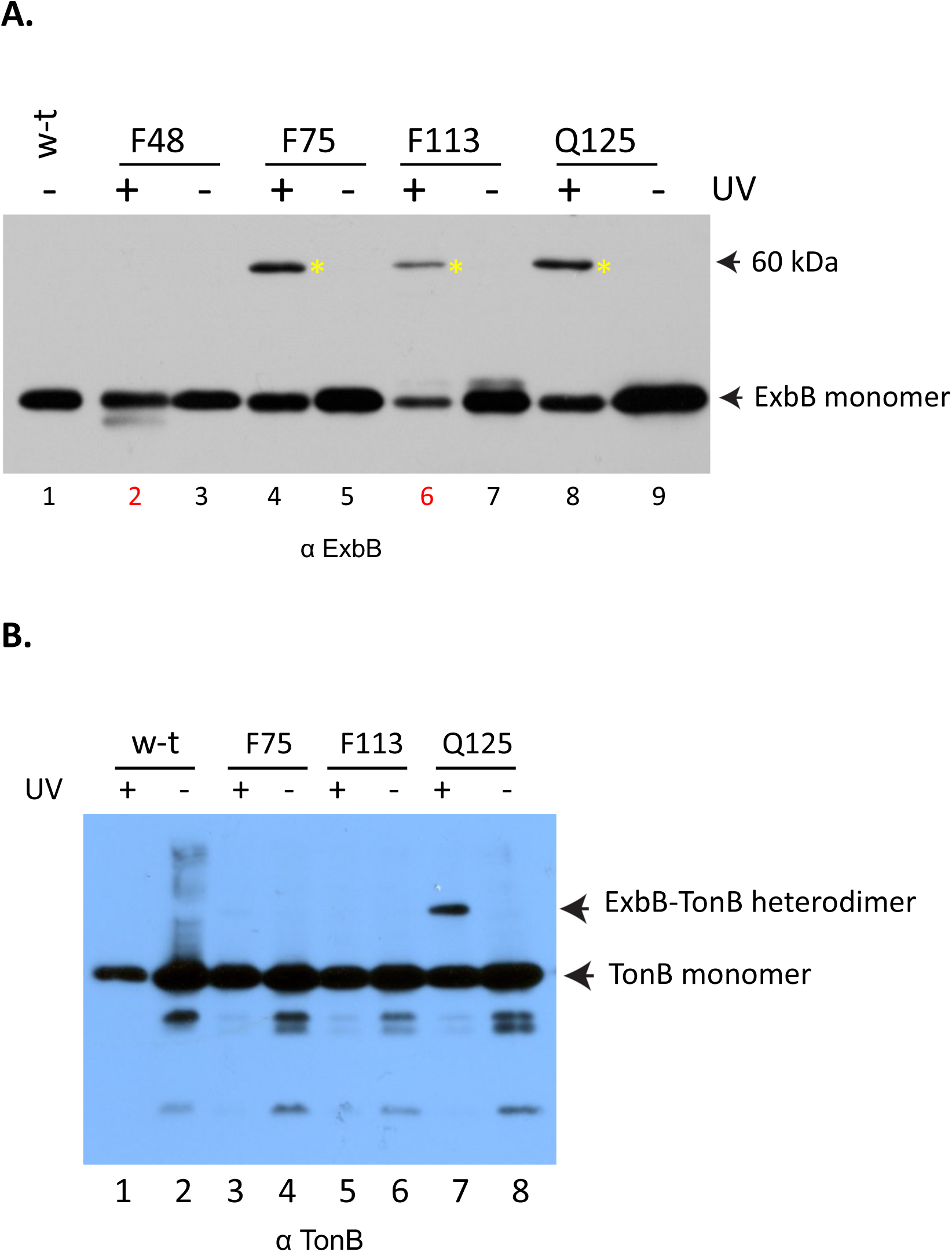
The ExbB cytoplasmic loop forms a ∼60 kDa complex with an unknown protein. Plasmid-encoded ExbB variants with pBpa substituted at residues F48, F75, F113, and Q125 in the cytoplasmic loop were photo-cross-linked *in vivo* with UV light (+) and under the same conditions in the absence of UV light (-). “w-t” stands for wild-type, W3110. The background strain for photo-cross-linking, RA1017 (W3110 Δ*exbB/D*, Δ*tolQRA*), also carried the pEvol plasmid (58). Equivalent numbers of exponential-phase bacteria were evaluated by immunoblotting. **A.) Immunoblot with anti-ExbB polyclonal antibodies.** The positions of ExbB monomer and the ∼60 kDa complex it forms (yellow asterisks) are indicated on the right. Lanes 2 and 6 labels (bottom) are red to indicate low recovery of sample indicated by subsequent Coomassie-staining of the immunoblot polyvinylidene difluoride (PVDF) membrane. **B)**. **Immunoblot with anti-TonB monoclonal 4F1 antibodies.** The positions of TonB monomer and the ExbB-TonB complex formed by ExbB Q125 (pBpa) are indicated on the right.

Although TonB has a calculated mass of 26.1 kDa, it has an aberrantly high apparent molecular mass of ∼36 kDa on SDS polyacrylamide gels due to its proline-rich region (62, 63). It was therefore a candidate to be part of the ∼60 kDa photo-cross-linked complex with ExbB. Immunoblot analysis of the *in vivo* cross-linking profiles of ExbB F78 (pBpa), F113 (pBpa) and Q125 (pBpa) with anti-TonB antibody revealed that only ExbB Q125 (pBpa) near the cytoplasmic side of transmembrane domain 2 (Fig. 1B) photo-cross-linked with TonB (Fig. 5B, lane 7). Thus, complexes formed by ExbB F78 (pBpa) and F113 (pBpa) that did not cross-link to TonB (Fig. 5B, lanes 3, 5) constituted cross-links to the ∼34 kDa unknown protein.

Consistent with those results, photo-cross-linking through cytoplasmic loop residue ExbB S80 (pBpa) also identified a ∼60 kDa complex that was not TonB, since it was also detected in a Δ*tonB* background (Fig. 6A, lanes 5, 7), and not ExbD, because no UV-specific ExbB complexes were detected by immunoblot with anti-ExbD antibodies (Fig. 6B). Since ExbB S80 is located between F75 and F113, it seemed possible that the ∼60 kDa complex consisted of the same unknown protein as the ∼60 kDa complex identified in Fig. 5A. ExbB S80 (pBpa) also formed a ∼52 kDa complex, almost certainly the previously identified ExbB formaldehyde cross-linked homodimer (38). The fact that ExbB S80 (pBpa) could be bound to two different proteins (a second ExbB and an unknown protein) was consistent with the proposed dynamics of a TonB-dependent energy transduction cycle (Fig. 2).

**Figure 6.**
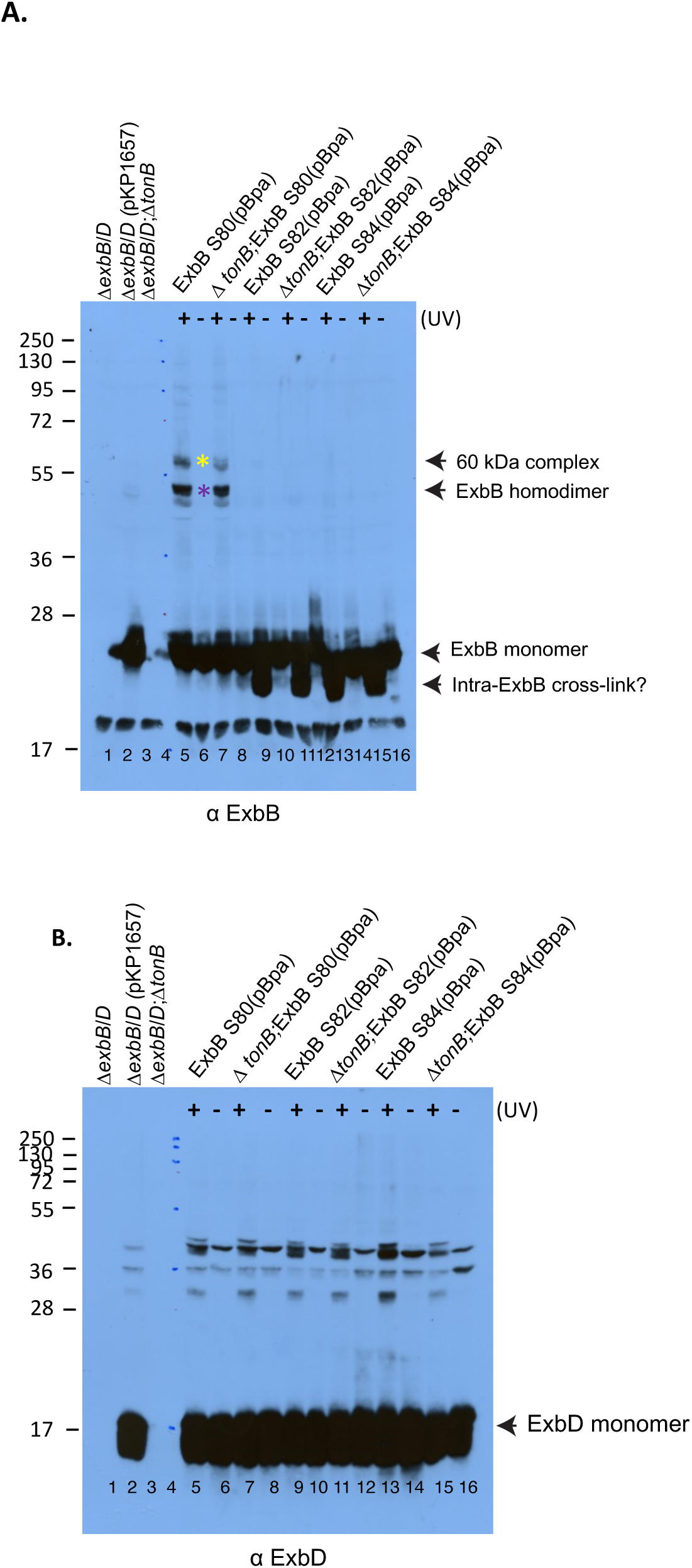
S80 (pBpa) in the ExbB cytoplasmic loop makes a ∼60 kDa complex with an unknown protein. Plasmid-encoded ExbB variants with pBpa substituted at residues S80, S82, and S84 were photo-cross-linked *in vivo* with UV light (+) and in the absence of UV light (-). Equivalent numbers of exponential-phase bacteria were evaluated by immunoblotting. Δ*exbB/D* stands for strain RA1017; Δ*exbB/D*, Δ*tonB* stands for strain KP1566, a derivative of RA1017 (see Table 4). pKP1657 is the parent plasmid expressing both ExbB and ExbD and the corresponding ExbB variants. All strains also contained plasmid pEvol (58). The positions of molecular mass standards in lane 4 are indicated on the left. **A.)** An immunoblot developed with anti-ExbB antibodies is shown. The positions of ExbB monomer, ExbB homodimer (purple asterisk), a ∼60 kDa complex containing both ExbB and an unknown protein (yellow asterisk), and a potentially intramolecularly cross-linked ExbB monomer are indicated on the right. **B.)** An immunoblot of the same samples from A.) developed with anti-ExbD antibodies. The position of ExbD monomer is indicated on the right.

ExbB S82 (pBpa) and S84 (pBpa) formed no cross-linked complexes, testifying to the specificity of the cross-links formed by S80 (pBpa). Both ExbB S82 (pBpa) and S84 (pBpa) gave rise to ExbB-specific bands of lower mass than the monomer (Fig. 6A, lanes 9, 11, 13, 15). Because the lower-mass bands were UV-specific, they were most likely the result of intra-protein cross-links known to decrease the apparent masses of proteins in SDS polyacrylamide gel electrophoresis (64). The finding of intra-protein cross-links was consistent with the hypothesized ability of ExbB cytoplasmic loops to protect the TonB amino terminus from labeling, depending on whether the TonB periplasmic domain was interacting with FepA (65).

As a bacterial actin homologue (41), the cytoplasmic protein MreB could serve to move tetrameric ExbB through an energy transduction cycle (Fig 2). The mass of MreB (37 kDa) also made it a candidate to be part of the UV-specific ∼60 kDa complex observed in Figs. 5A and 6A. However, none of the complexes contained MreB based on immunoblot analysis with anti-MreB antiserum (data not shown). In addition, the compound A22 [S-(3,4-dichlorobenzyl) isothiourea], a known inhibitor of MreB (66), had no effects on S80 pBpa formation of either complex (data not shown).

Taken together, the *in vivo* photo-cross-linking data showed that ExbB formed a ∼60 kDa complex through multiple pBpa substitutions in the ExbB cytoplasmic loop, and that the other protein in the complex was not MreB (37 kDa), not TonB (∼36 kDa), not a second ExbB (∼26 kDa) and not ExbD (∼17 kDa). Thus, the cytoplasmic loop of ExbB interacted with an unknown cytoplasmic protein of ∼34 kDa.

### The ExbB periplasmic loop

The periplasmic loop of ExbB that connects transmembrane domains 2 and 3 in Fig. 1B extends from residues ∼153-177 (31). We substituted the segment from 156-161 (FIGIAQ) with a block of six Alanine residues in plasmid pKP1766 (Table 4) and assayed its ability to complement strain RA1017 (W3110 Δ*exbBD*::*kan*, Δ*tolQRA*) in an assay of initial iron transport rates. Rather than FIGIAQ simply being a place holder between two transmembrane domains, ExbB (156-161 Ala) appeared to have an important function, since it was entirely inactive for iron transport (data not shown). Consistent with that result, it did not complement RA1017 with respect to colicins B and Ia sensitivity. Because an Alanine residue scan of the individual residues 156-161 did not identify a single residue that knocked out activity, it seemed likely that two or more residues were involved (Kopp and Postle, unpublished results).

### The ExbB cytoplasmic carboxy terminus

#### ExbB Q215-L224 contain the key residues in the ExbB cytoplasmic carboxy terminus

Deletion of the cytoplasmic carboxy terminal tail (residues 195-244, Fig. 4B) renders ExbB inactive (38). To characterize it more fully, we engineered five sequential 10-residue deletions (Δ10s) and a corresponding set of five 10-residue Alanine-substituted regions (10-Alas). Each of the variants was initially characterized as to level of arabinose inducer required to achieve chromosomally encoded levels of expression from the arabinose promoter in the pBAD24 backbone of pKP660 as well as the ability to support initial rates of iron transport under those conditions (Table 2).

**Table 2.** Comparison of Δ10 and 10-Ala variants in the ExbB cytoplasmic tail.

|  | % Arabinose for<br>Chromosomal Expression<sup>a</sup> |  | Activity<sup>b</sup> |  |
| --- | --- | --- | --- | --- |
| | $\Delta 10$ | 10-Ala | $\Delta 10$ | 10-Ala |
| Wild-type<sup>c</sup> | 0.00025% | 0.0005% | 100% | 100% |
| 195-204<sup>d</sup> | 0.2% | 0.0001% | 0% | 0% |
| 205-214 | 0.2% | 0.0005% | 0% | 50% |
| 215-224 | 0.2% | 0.2% | 0% | 0% |
| 225-234 | 0% | 0% | 100% | 140% |
| 235-244 | 0% | 0.0001% | 35% | 60% |

<sup>a</sup>An indirect measure of sensitivity to endogenous proteases. The higher the level of arabinose required to achieve chromosomally encoded levels of expression, the greater the proteolytic sensitivity.
<sup>b</sup>Activity measured as a % of initial rates of iron transport compared to wild-type W3110. Effects of the $\Delta 10$ and 10-Ala substitutions were measured in the RA1017 strain background (W3110 $\Delta exbBD::kan$ , $\Delta tolQRA$ ). ExbB 195-204 Ala is inactive due to presence of the Y195A substitution (38).
<sup>c</sup>Wild-type is defined as plasmid pKP660 in RA1017.
<sup>d</sup>Numbers refer to the *exbB* codons that were modified in plasmid pKP660.

The three ExbB Δ10s closest to the transmembrane domain 3 (ΔY195-G204, ΔF205-A214 and ΔQ215-L224) required the maximally effective level of 0.2% (w/v) L-arabinose to achieve near chromosomal level of expression, indicating that they were proteolytically unstable and suggesting that the structural perturbation they caused engaged endogenous proteases (Table 2). In contrast the two distal Δ10s (ΔD225-P234 and ΔV235-G244) were proteolytically stable since 0% arabinose was required for them to be expressed at chromosomal levels (Table 2). The three ExbB Δ10s closest to the transmembrane domain 3 were also inactive whereas the distal two Δ10s retained activity in assays of initial rates of iron transport (Table 2) and sensitivities to colicin B (data not shown) under conditions where all ExbB variants were expressed at chromosomal levels. These results mirrored our previous ones from the ExbB cytoplasmic domain Δ10s (Table 1), where the only proteolytically stable Δ10 is also the only one that retains some activity (ExbB Δ100-109).

Among the five ExbB 10-Alas, only ExbB 215-224 Ala required 0.2% (w/v) L-arabinose to achieve chromosomally encoded levels of expression; it was therefore judged to be proteolytically unstable (Table 2). ExbB Q215-L224 Ala also appeared to be the only 10-Ala substitution where inactivity was due to an unexplained loss of important residues; the inactivity of ExbB Y195-G204 Ala was almost certainly due to the previously identified inactivity of ExbB Y195A (38) while ExbB F205-A214 Ala, D225-P234 Ala, and V235-G244 Ala were at least 50% active (Table 2).

Based on our study of the ExbB cytoplasmic loop (28), comparison of immediate growth arrest phenotypes for the cytoplasmic tail variants identified ExbB Q215-L224 as potential region of interaction with an unknown protein (Fig. 4B). Of the ExbB Δ10s, only the three most proximal to the transmembrane domain 3, encompassing residues 195-224, exhibited immediate growth arrest. Phenotypes of the 10-Ala substitutions in the cytoplasmic tail narrowed the focus to ExbB Q215-L224 Ala which was the only one of the five 10-Alas that also exhibited immediate growth arrest (Table 3; Figs. 4B, 7). This result was unexpected since none of the cytoplasmic loop 10-Alas exhibited any immediate growth arrest [Table 1, Fig. 4A].

**Figure 7.**
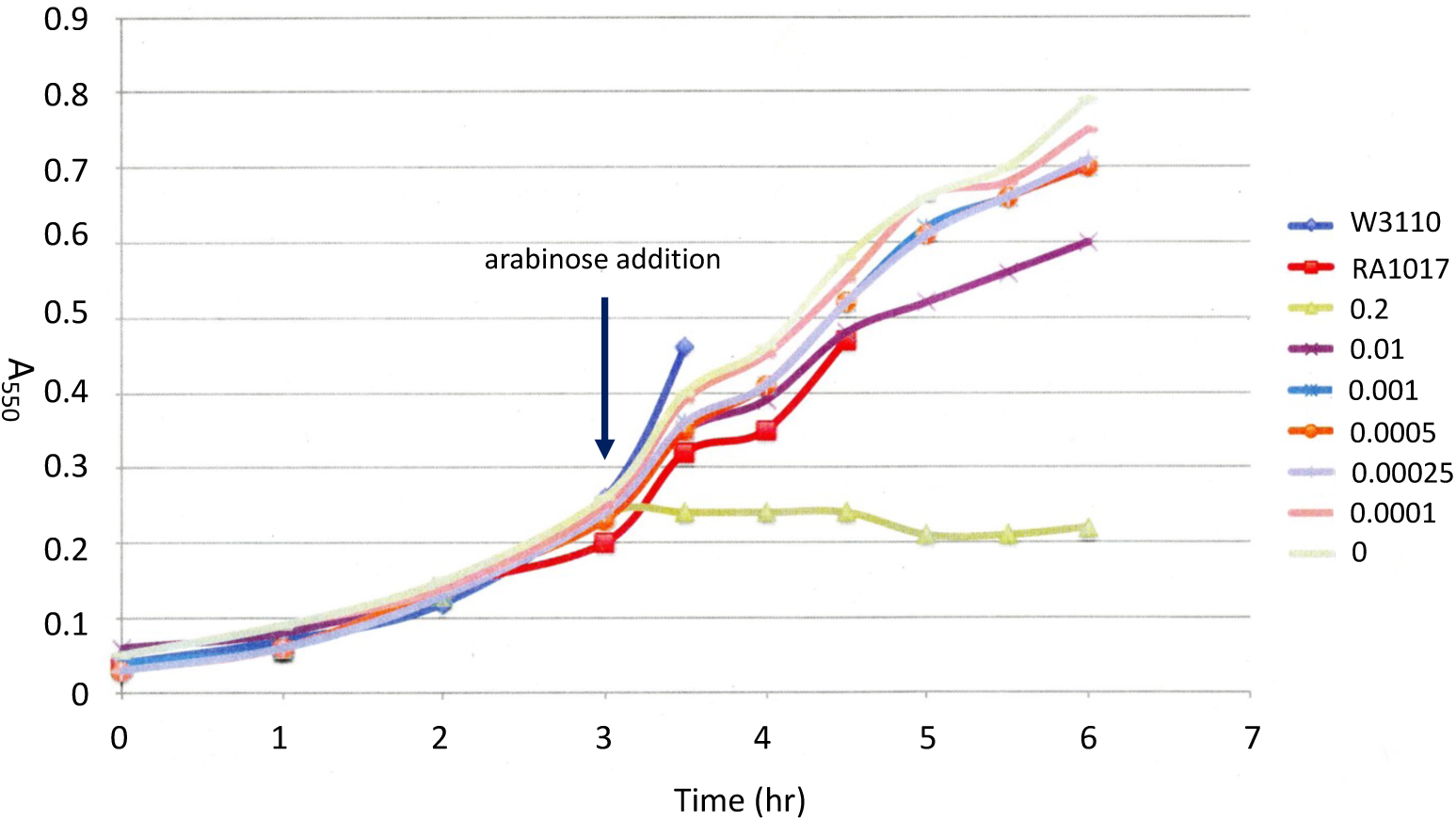
Maximal induction of ExbB 215-224 Ala expression with 0.2% arabinose results in immediate growth arrest. Subcultures of strain RA1017 carrying pKP2009 (ExbB 215-224 Ala) were grown to mid-logarithmic phase in M9 medium with 100 µg/ml ampicillin. Three hours after inoculation, the various percentages of arabinose listed to the right of the graph were distributed among the subcultures and the A_550_ monitored every 30 min (left). The time scale is shown at the bottom. Growth of W3110 and RA1017 were monitored as controls without added arabinose.

**Table 3.** Effect of 0.2% arabinose on growth of ExbB cytoplasmic tail variants^a^.

| Plasmid | initial A<sub>550</sub> | A<sub>550</sub> after 1 hr |
| --- | --- | --- |
| pKP660 (wild-type ExbB/D) | 0.25 | 0.50 |
| pKP1963 (ExbB Δ195-204) | 0.23 | 0.22 |
| pKP1964 (ExbB Δ205-214) | 0.24 | 0.22 |
| pKP1965 (ExbB Δ215-224) | 0.24 | 0.25 |
| pKP1940 (ExbB Δ225-234) | 0.28 | 0.48 |
| pKP1912 (ExbB Δ235-244) | 0.28 | 0.47 |
| pKP2019 (ExbB 195-204 Ala) | 0.17 | 0.39 |
| pKP2008 (ExbB 205-214 Ala) | 0.12 | 0.31 |
| pKP2009 (ExbB 215-224 Ala) | 0.24 | 0.22 |
| pKP2025 (ExbB 225-234 Ala) | 0.24 | 0.39 |
| pKP2010 (ExbB 235-244 Ala) | 0.31 | 0.50 |

<sup>a</sup>Plasmids expressing ExbB variants were assayed by subculturing from saturated LB cultures and allowing them to grow in T-broth without inducer to ~ A<sub>550</sub> = 0.24 (initial A<sub>550</sub>), adding 0.2% arabinose, and assessing the A<sub>550</sub> again after 1 hour. Numbers in red indicate ExbB variants that exhibited immediate growth arrest.

#### The ExbB Q215-D223 region interacts with unknown proteins of ∼ 24 kDa and ∼29 kDa

To test the idea that residues Q215-D223 (L224 was not investigated in this study) could be a site of interaction with unknown proteins, they were each submitted to an *in vivo* photo-cross-linking analysis. In the presence of UV light, ExbB Q215 (pBpa) formed ExbB homodimers (Fig. 8A, lanes 4, 6), seen previously with formaldehyde cross-linking (38) and that were absent from a corresponding immunoblot developed with anti-ExbD antibodies (Fig. 8B, lanes 4, 6).

**Figure 8.**
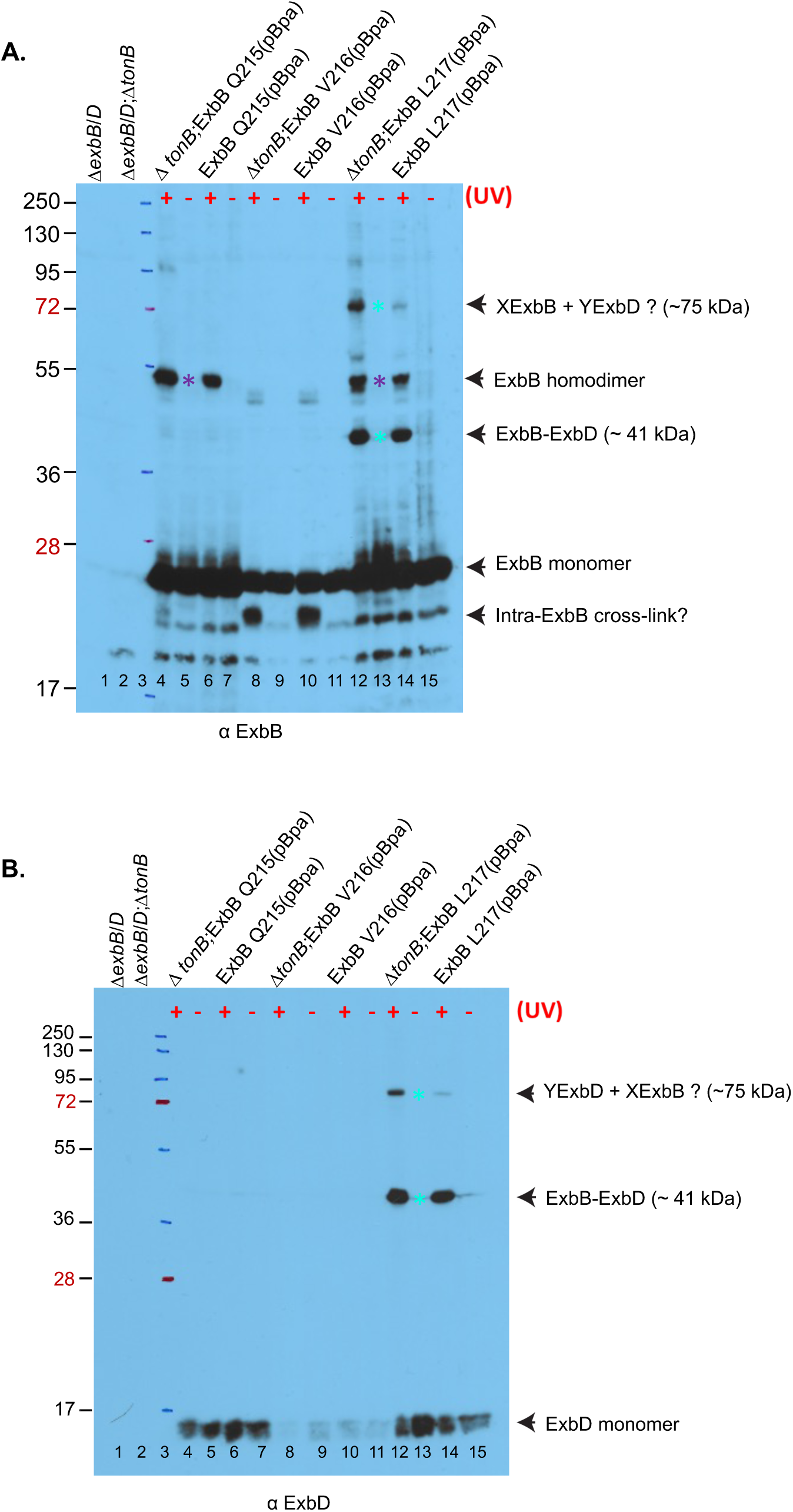
Photo-cross-linking through ExbB Q215, V216, L217 identifies known ExbB-ExbD interactions. Plasmid-encoded ExbB variants with pBpa substituted at residues Q215, V216, and L217 were photo-cross-linked *in vivo* with UV light (+) and under the same conditions in the absence of UV light (-). Equivalent numbers of exponential-phase bacteria were evaluated by immunoblotting. Δ*exbB/D* stands for strain RA1017; Δ*exbB/D*, Δ*tonB* stands for strain KP1566, a derivative of RA1017 (see Table 4). pKP1657 is the parent plasmid expressing both ExbB and ExbD and the corresponding ExbB variants. All strains also contained plasmid pEvol (58). The positions of molecular mass standards in lane 3 are indicated on the left. **A.)** An immunoblot developed with anti-ExbB antibodies is shown. The positions of ExbB monomer, the ExbB homodimer (purple asterisk), the ExbB-ExbD complex (aqua asterisk) and a potentially intramolecularly cross-linked ExbB monomer are indicated on the right. **B)** An immunoblot of the samples from A.) developed with anti-ExbD antibodies is shown. The positions of ExbD monomer and the ExbB-ExbD complex (aqua asterisk) are indicated on the right.

Complexes formed by ExbB V216 (pBpa) were difficult to evaluate since they appeared to be proteolytically unstable under the photo-cross-linking conditions (Fig. 8A, lanes 8-11)—a result confirmed by the decreased expression of ExbD in the corresponding samples (Fig. 8B, lanes 8-11), since ExbD relies on ExbB for proteolytic stability (24). In Fig. 8A, lanes 8 and 10, ExbB V216 (pBpa) gave rise to apparently intramolecular UV-specific ExbB bands with apparent masses smaller than the monomer, like those seen for ExbB S82 (pBpa) and ExbB S84 (pBpa) (Fig. 6A, lanes 9, 11, 13, 15).

ExbB L217 (pBpa) formed UV-specific ExbB homodimers and a heterodimer consisting of one ExbB and one ExbD at ∼41 kDa that has also been captured with formaldehyde cross-linking (19). A still larger heterodimer at ∼75 kDa must also have contained at least one ExbB and one ExbD (compare Fig. 8A, lanes 12, 14 and Fig. 8B, lanes 12, 14). The absence of TonB increased the abundance of this complex, possibly because TonB was no longer present to compete with ExbD for ExbB interactions. TonB formaldehyde cross-links individually to both ExbB and ExbD *in vivo*, with the TonB-ExbD complex being PMF-dependent (19, 27).

It was not possible to identify with confidence the number of components in the ∼75 kDa XExbB-YExbD complex, based on its mass (Fig. 8). Two ExbB (26 kDa) and one ExbD (15 kDa) would total only ∼67 kDa and would require one ExbB (pBpa) to photo-cross-link to a second ExbB (pBpa), and the second ExbB pBpa to cross-link to ExbD. A complex of three ExbB (pBpa) plus one ExbD would have an apparent mass of 93 kDa, suggesting it was also not representative of the ∼75 kDa complex. A complex of two ExbB (pBpa) plus two ExbD, which best approximates the observed ∼75 kDa complex, should presumably fall apart as it was boiled in SDS-containing sample buffer prior to electrophoresis. However, in an experiment described below, an ExbB/ExbD complex appeared to be resistant to denaturation under those circumstances, suggesting that an ExbB_2_/ExbD_2_ complex was possible. Alternatively, the 75 kDa XExbB-YExbD complex reflected the unusual conformation that formaldehyde cross-linked ExbB tetramer + X can form, causing its apparent mass to vary with the percentage of cross-linking in the gel matrix (67).

ExbB L218 (pBpa) formed UV-specific ExbB homodimers, ExbB-ExbD heterodimers and three larger ExbB- and UV-specific complexes (Fig. 9, lanes 4, 6; purple, aqua, and yellow asterisks respectively). The three larger complexes could include ExbD or unknown proteins, but to account for the larger mass, likely included multiple ExbBs. The largest complex might be the ExbB tetramer + X (38). Unfortunately, we were unable to obtain the corresponding anti-ExbD immunoblot for Fig. 9 before the COVID-19 pandemic emerged, the corresponding author retired, and the lab was permanently closed.

**Figure 9.**
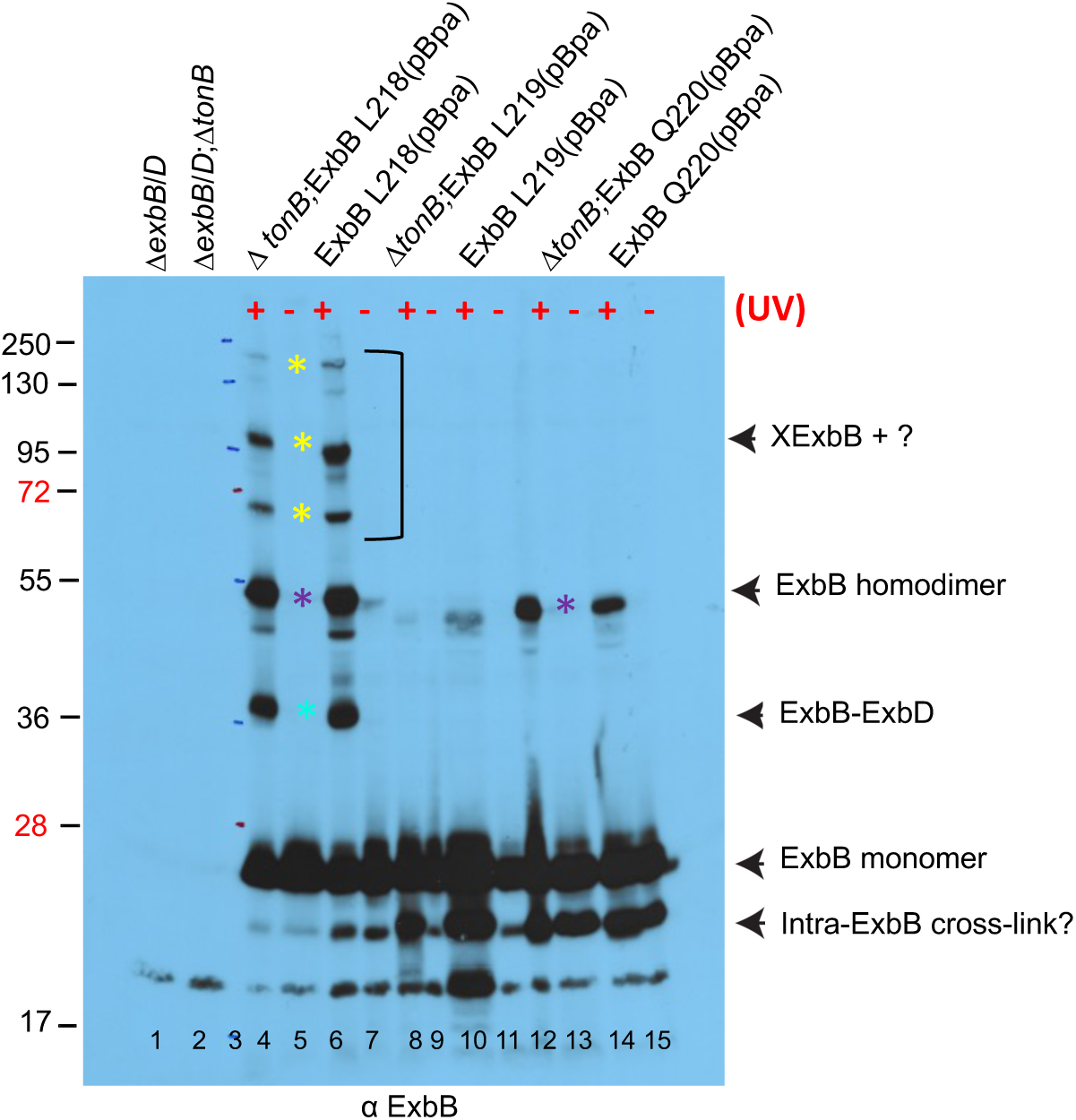
Photo-cross-linking through ExbB L218, L219, Q220 identifies complexes with known and potentially, with unknown proteins. Plasmid-encoded ExbB variants with pBpa substituted at residues L218, L219, and Q220 were photo-cross-linked *in vivo* with UV light (+) and under the same conditions in the absence of UV light (-). Equivalent numbers of exponential-phase bacteria were evaluated by immunoblotting. Δ*exbB/D* stands for strain RA1017; Δ*exbB/D*, Δ*tonB* stands for strain KP1566, a derivative of RA1017 (see Table 4). pKP1657 is the parent plasmid expressing both ExbB and ExbD and the corresponding ExbB variants. All strains also contained plasmid pEvol (58). An immunoblot developed with anti-ExbB antibodies is shown. The positions of ExbB monomer, the ExbB homodimer (purple asterisk), the ExbB-ExbD complex (aqua asterisk) and ExbB-specific complexes of unknown composition (yellow asterisk) are indicated on the right. The positions of molecular mass standards in lane 3 are indicated on the left.

ExbB L219 (pBpa) appeared to form the UV-specific intra-protein cross-link seen with ExbB S82 (pBpa) and ExbB S84 (pBpa), but no significant level of higher mass complexes (Fig. 9, lanes 8, 10). ExbB Q220 (pBpa) formed UV-specific homodimers (Fig. 9, lanes 12, 14). While it also gave rise to smaller mass ExbB-specific bands (lanes 12-15), they did not appear to be UV-specific and were therefore likely to be proteolytic degradation products.

ExbB S221, R222, and D223 (pBpa) derivatives formed the UV-specific ExbB homodimer at ∼52 kDa (Fig. 10, lanes 4, 8, 12, purple asterisks). The ExbB S221 (pBpa) homodimer formation was not as abundant as that seen with ExbB R222 and D223. It appeared to be TonB-dependent, suggesting that the conformation of ExbB *in vivo* differed depending on the presence or absence of TonB. In addition, ExbB S221 (pBpa) formed an abundant ∼41 kDa ExbB-ExbD complex (Fig. 10A, aqua asterisk, lanes 4, 6 and Fig. 10B, lanes 4, 6). ExbB S221, R222, and D223 (pBpa) all appeared to be subject to endogenous proteolysis (Fig. 10, lanes 4-15).

**Figure 10.**
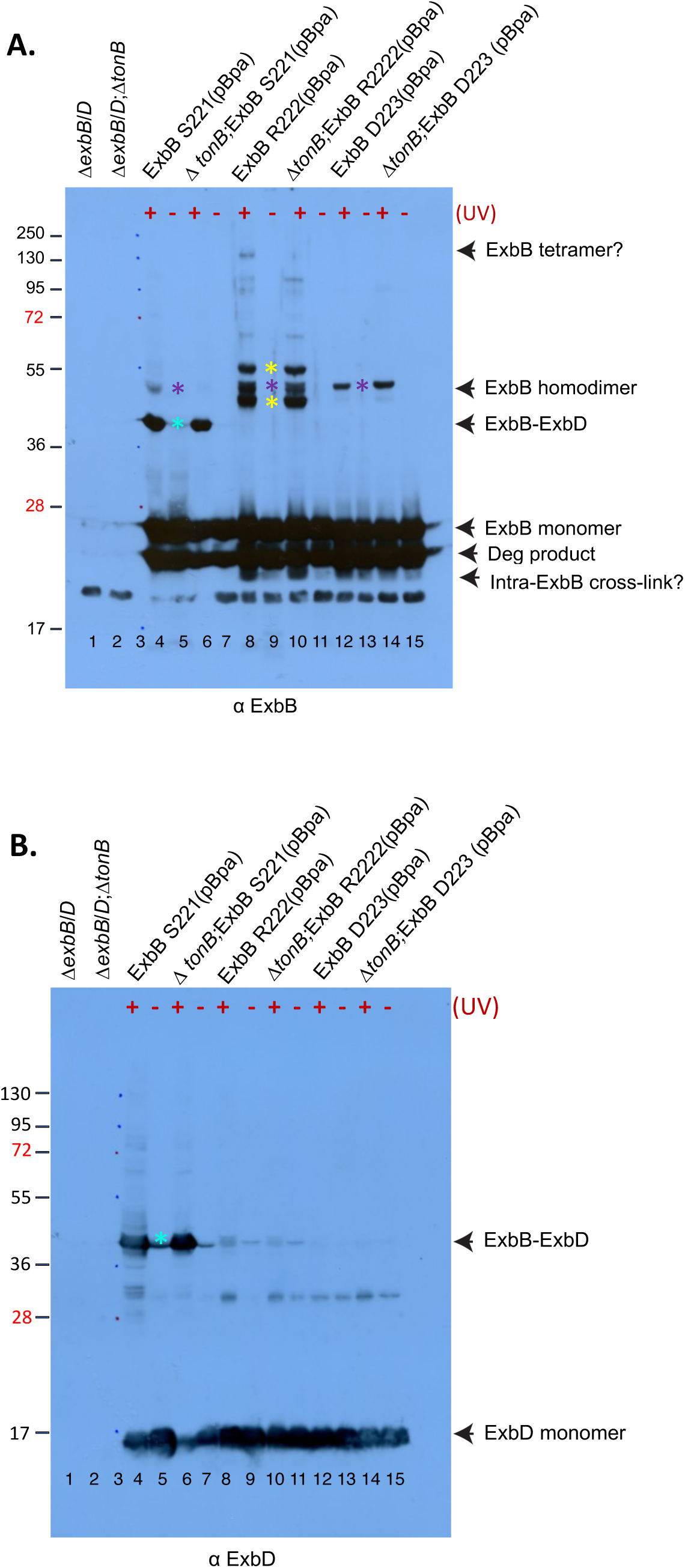
Photo-cross-linking through ExbB S221, R222, and D223 identifies complexes with known and unknown proteins. Plasmid-encoded ExbB variants with pBpa substituted at residues S221, R222, and D223 were photo-cross-linked *in vivo* with UV light (+) and under the same conditions in the absence of UV light (-). Equivalent numbers of exponential-phase bacteria were evaluated by immunoblotting. Δ*exbB/D* stands for strain RA1017; Δ*exbB/D*, Δ*tonB* stands for strain KP1566, a derivative of RA1017 (see Table 4). pKP1657 is the parent plasmid expressing both ExbB and ExbD and the corresponding ExbB variants. All strains also contained plasmid pEvol (58). The positions of molecular mass standards in lane 3 are indicated on the left. **A.)** An immunoblot developed with anti-ExbB antibodies is shown. The positions of ExbB monomer, the ExbB homodimer (purple asterisk), the ExbB-ExbD complex (aqua asterisk), degradation products and likely intramolecularly cross-linked monomer are indicated on the right. The yellow asterisk identifies ExbB complexes with unknown proteins. **B)** An immunoblot of the samples from A.) developed with anti-ExbD antibodies is shown. The positions of ExbD monomer and the ExbB-ExbD complex (aqua asterisk) are indicated on the right.

ExbB R222 is notable as the most highly conserved member of a conserved S_221_RDL_224_ motif (Postle and Kippelen, data not shown). In addition to forming a homodimer, ExbB R222 (pBpa) also formed abundant complexes with two unknown proteins (Fig. 10A, yellow asterisk, lanes 8, 10): 1) a ∼55 kDa complex suggesting an unknown protein of ∼29 kDa; and 2) a ∼50 kDa complex suggesting an unknown protein of ∼24 kDa. A significantly less abundant series of higher mass complexes were also observed. Of these, the ∼130 kDa complex was notable because it appeared to be TonB-dependent (Fig. 10A, lanes 8, 10). Based on size, the ∼130 kDa complex could be an ExbB tetramer (38) that required the presence of TonB to form. It did not appear to include any ExbD (Fig. 10B, lanes 8, 10). ExbB R222 (pBpa) also appeared to form UV-specific intra-protein cross-links (Fig. 10, lanes 8-11). Although we consider it to be unlikely, the possibility that the ∼24 kDa protein was a proteolytic product of the ∼29 kDa protein was not ruled out.

### Studies on TonB transmembrane domain interactions

#### The TonB transmembrane domain photo-cross-links to a second TonB in vivo

TonB can be functionally divided into two domains. The TonB amino terminus contains an uncleaved signal anchor (residues 12-31) essential for export and activity (48, 68), while the majority of TonB occupies the periplasmic space (Fig. 1).

In this study, the ability of TonB to homodimerize through its core transmembrane domain was assessed in strain KP1566 (W3110, Δ*tonB*, Δ*exbB/D*, Δ*tolQRA;* Table 4) which lacks any known chromosomally encoded interaction partners. Under these circumstances, plasmid-encoded TonB A22-, V23-, V24-(pBpa) each formed a UV-specific, TonB-specific homodimer at ∼68 kDa (Fig. 11, lanes 3, 5, 7). TonB A25 (pBpa) in lanes 9 and 10 appears to be proteolytically unstable, making formation of homodimers difficult to assess.

**Figure 11.**
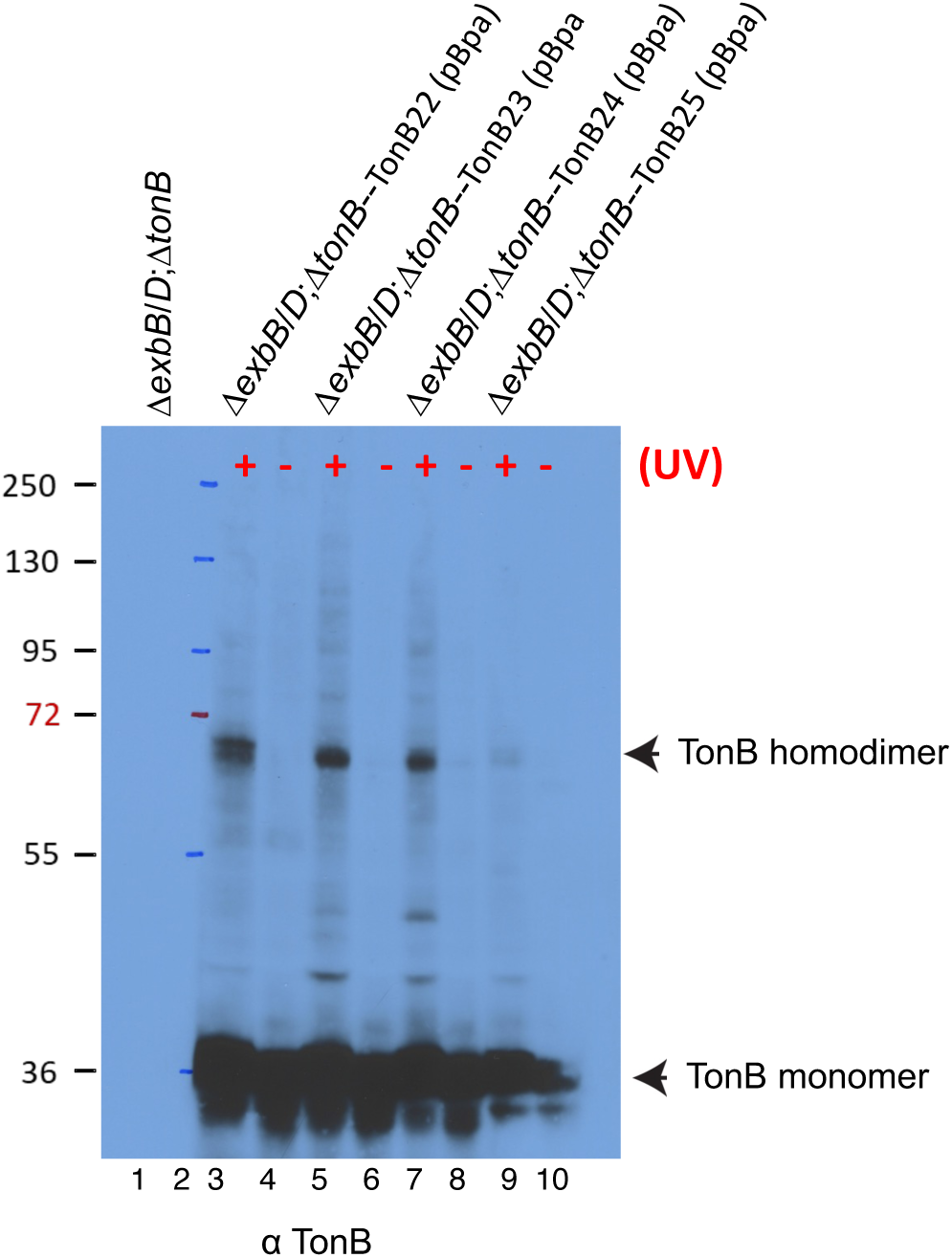
The TonB transmembrane domain photo-cross-links to a second TonB. Plasmid-encoded TonB variants with pBpa substituted at residues A22, V23, V24, and Ala25 were photo-cross-linked *in vivo* with UV light (+) and under the same conditions in the absence of UV light (-). Equivalent numbers of exponential-phase bacteria were evaluated by immunoblotting. Δ*exbB/D*;Δ*tonB* stands for strain KP1566, a *tonB* derivative of RA1017. All strains also contained plasmid pEvol (58). An immunoblot developed with anti-TonB antibodies is shown. The TonB monomer and homodimer are indicated on the right. Positions of molecular mass standards seen in lane 2 are indicated on the left.

**Table 4.**
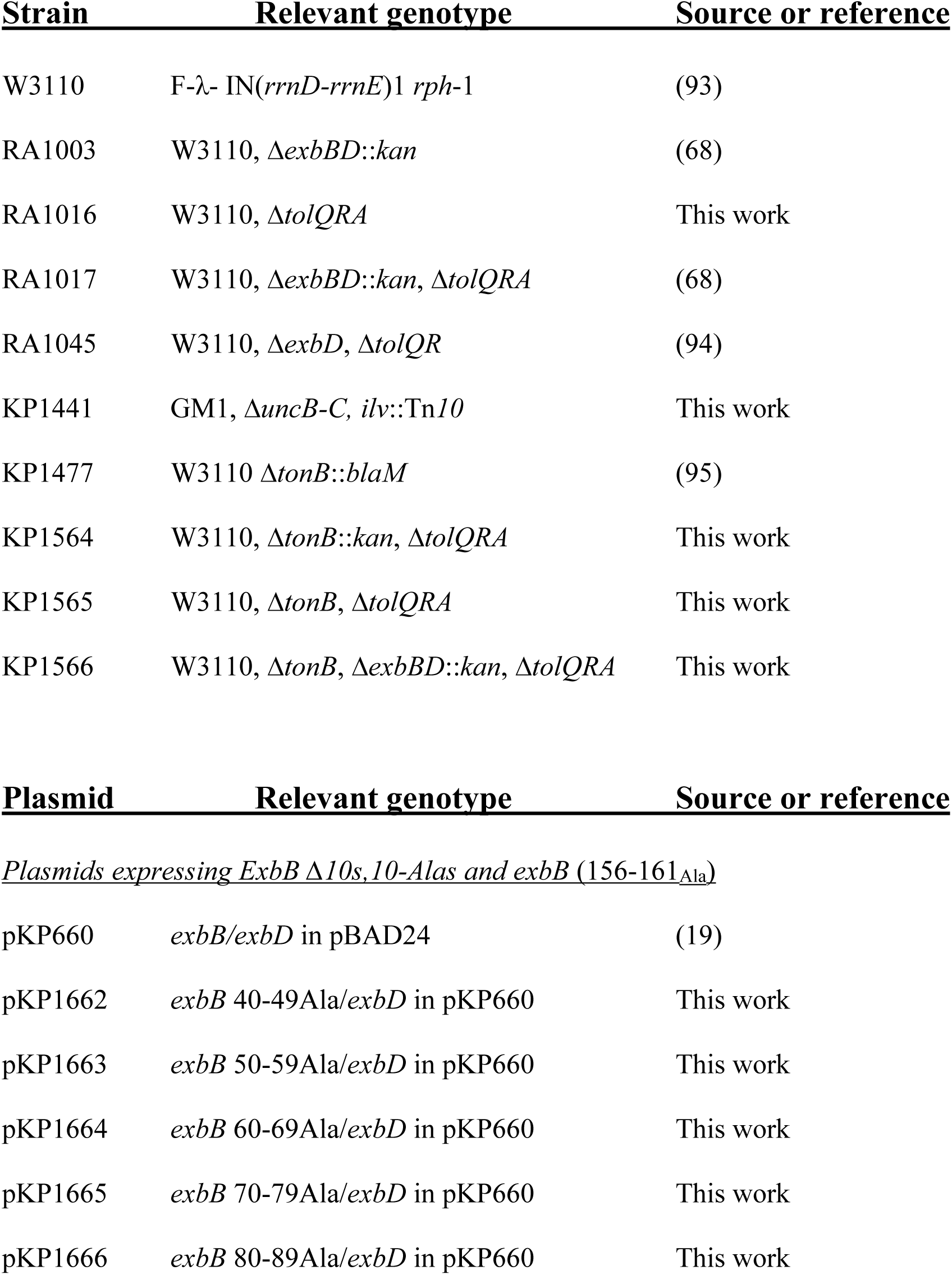
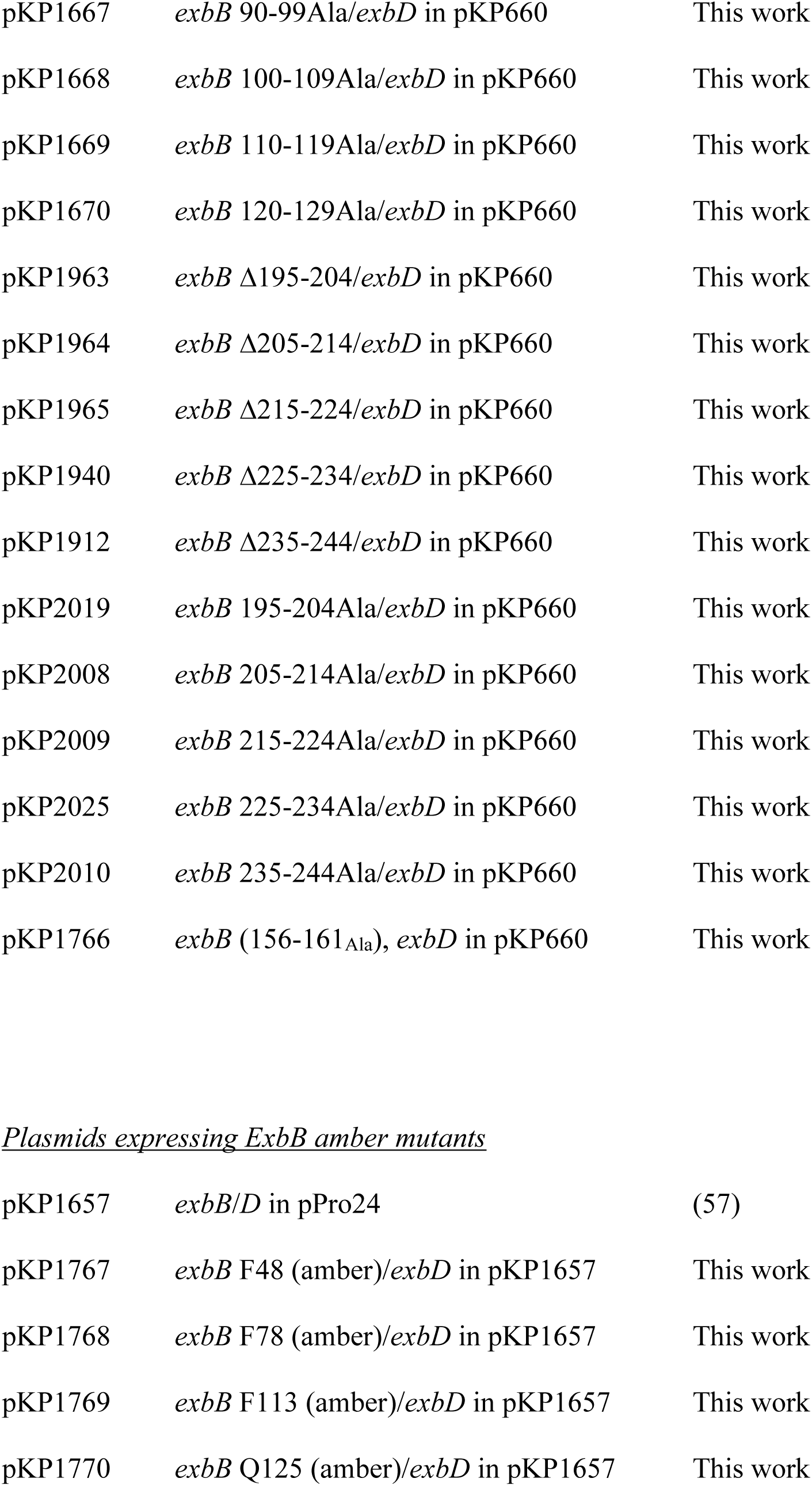
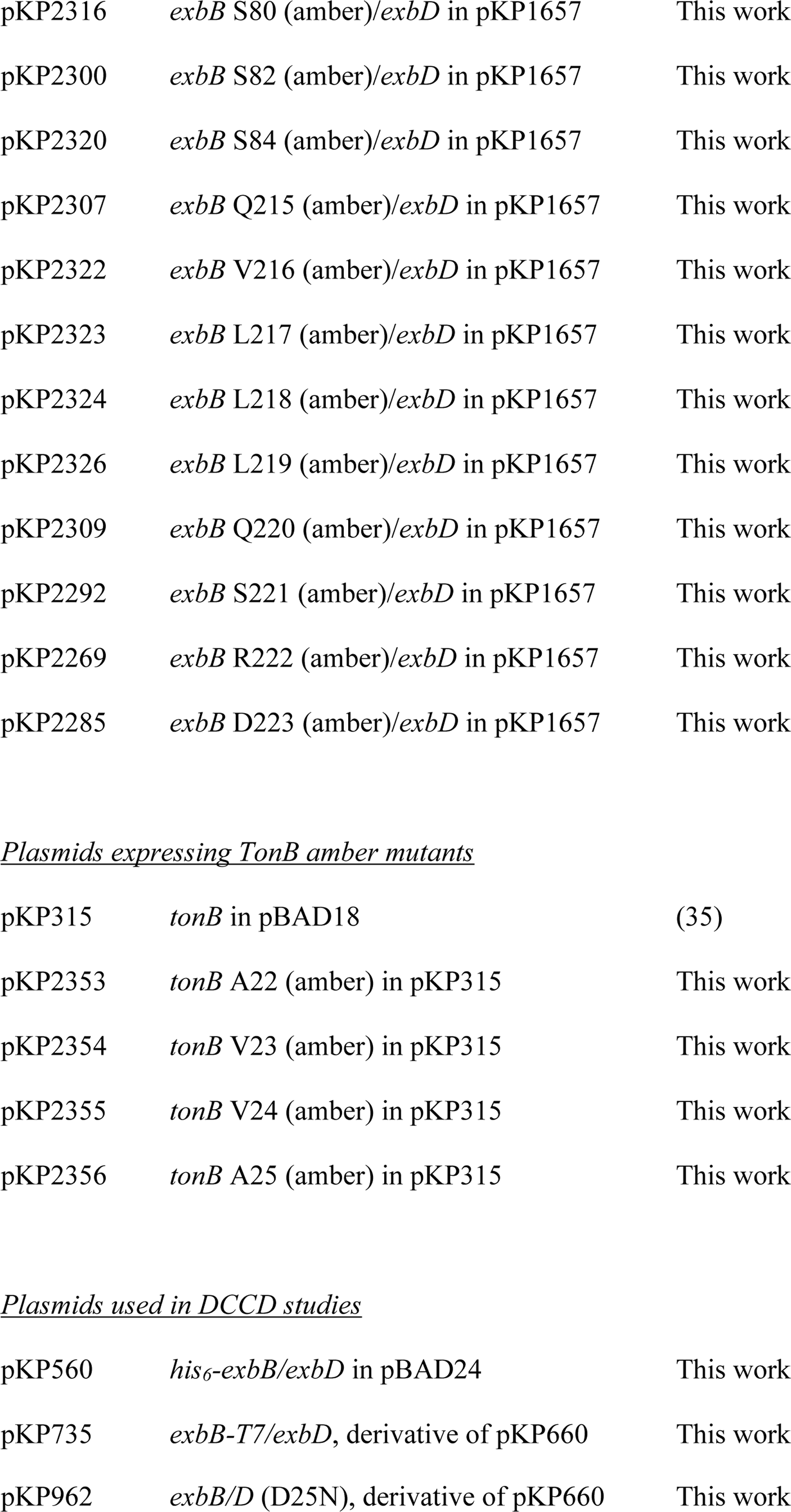
Strains and Plasmids used in this study.

Trapping of homodimers by three adjacent TonB pBpa substitutions suggested that the orientation of the transmembrane domains relative to one another changed during an energy transduction cycle (69). While that idea appeared to be contradictory to our previous observations where TonB P41C forms disulfide-linked homodimers that are functional *in vivo,* it may be that TonB dimers trapped through P41C disulfides still provide enough steric flexibility that the TonB transmembrane domains can move as needed [Fig. 2; (42)].

#### The TonB transmembrane domain also photo-cross-links to ExbD in vivo

ExbD has the same topology and uncleaved signal anchor configuration as TonB (Fig. 1). TonB and ExbD interact cyclically through their periplasmic domains, detected *in vivo* as PMF-dependent formaldehyde cross-linked heterodimers and disulfide-linked heterodimers [Fig. 2;(19, 42, 51, 52)].

Our previous results showed that I24 (pBpa) in the ExbD transmembrane domain forms heterodimers with TonB but did not identify the domain of TonB with which it interacted (57). To determine if the TonB transmembrane domain interacted directly with ExbD, the same set of plasmid-encoded TonB (pBpa) derivatives from Fig. 11 was photo-cross-linked in KP1477 (W3110 Δ*tonB*::*blaM*), immunoblotted, and analyzed with anti-ExbD antiserum (Fig. 12). In lanes 3, 5 and 9, a single UV-specific complex appeared at ∼45 kDa, an apparent mass that was significantly lower than that seen previously for formaldehyde cross-linked TonB-ExbD [∼52 kDa; (19)]. Because this ∼45 kDa complex was detected with specific anti-ExbD antibodies (6), was not detected in the control lacking the pEvol plasmids but expressing TonB pBpa derivatives (Fig. 12, lane 1) and could only have occurred by cross-linking with TonB (pBpa)s, we conclude that the TonB and ExbD transmembrane domains were within 3 Å of one another at some point in the energy transduction cycle (59). The ∼45 kDa complex’s small apparent mass may have been due to the sites through which the two proteins were connected (64).

**Figure 12.**
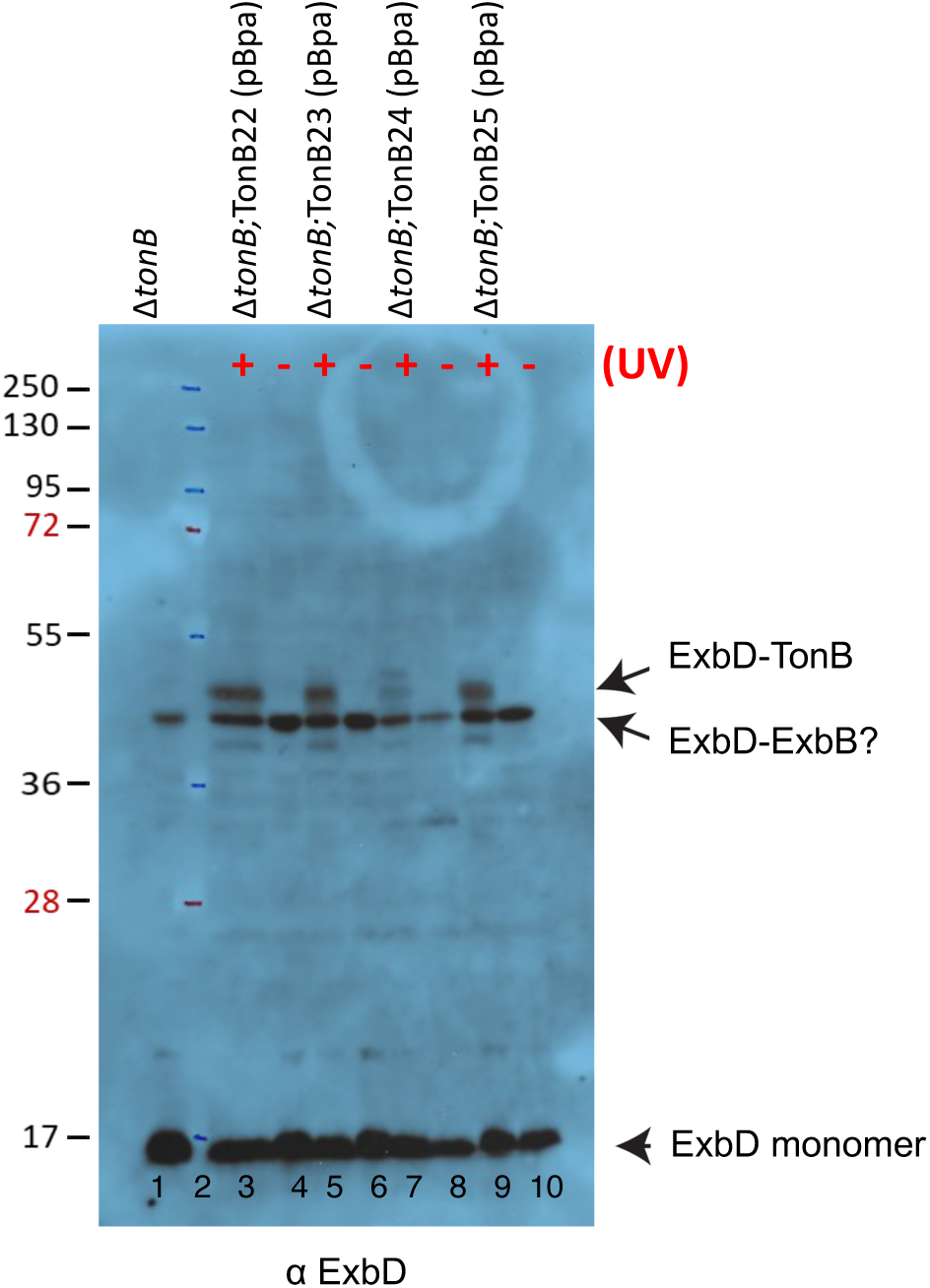
The TonB transmembrane domain photo-cross-links to ExbD. Plasmid-encoded TonB variants with pBpa substituted at residues A22, V23, V24, and Ala25 were photo-cross-linked *in vivo* with UV light (+) and under the same conditions in the absence of UV light (-). Equivalent numbers of exponential-phase bacteria were evaluated by immunoblotting. Δ*tonB* stands for strain KP1477 (W3110, Δ*tonB*, P14::*kan*) which expresses both ExbB and ExbD. All strains also contained plasmid pEvol (58). An immunoblot developed with anti-ExbD antibodies is shown. ExbD monomer, the TonB-ExbD complex and a likely ExbB-ExbD complex are indicated on the right. Positions of molecular mass standards seen in lane 2 are indicated on the left.

Also seen in Fig. 12 in control lane 1 and lanes 3-10 was an ExbD-specific complex at 41 kDa, the same mass seen when ExbD cross-links to ExbB (19). This complex appeared to be sufficiently strong that it tolerated sample preparation for gel electrophoresis since it occurred even in the absence of UV treatment.

### A new assay for ExbD dynamics: the effect of N, N_1_-Dicyclohexylcarbodiimide (DCCD)

Upon the discovery that unknown proteins bound to ExbB cytoplasmic domains and could thus be candidates to play a role in movement during energy transduction cycle or in immediate growth arrest, we wanted to determine if ATP synthase was also involved. If it was, the TonB system would join other *E. coli* membrane-dependent phenomena, such as protein export, that use both PMF and ATP as energy sources.

However, we found no difference in sensitivity to various colicins and phage between strains with and without ATP synthase (Δ*uncBC*—KP1441; Baker and Postle, unpublished). During that investigation, we also tested the *in vivo* effects of DCCD, a known inhibitor of ATP synthase in *E. coli* (70, 71). DCCD also inhibits other cellular processes including protein translocation into membrane vesicles and iron-sulfur reactions (72, 73) and can serve as a zero-length cross-linker (74). Consistent with the phage and colicin sensitivity assays, neither the absence of ATP synthase nor the presence of DCCD had any effect on previously characterized formaldehyde cross-linking profiles of TonB system proteins [(19, 57,61); data not shown]. However, DCCD unexpectedly decreased the initial rate of iron transport to ∼40% of the wild-type transport level, suggesting that it might modify individual TonB system proteins (Fig. 13).

**Figure 13.**
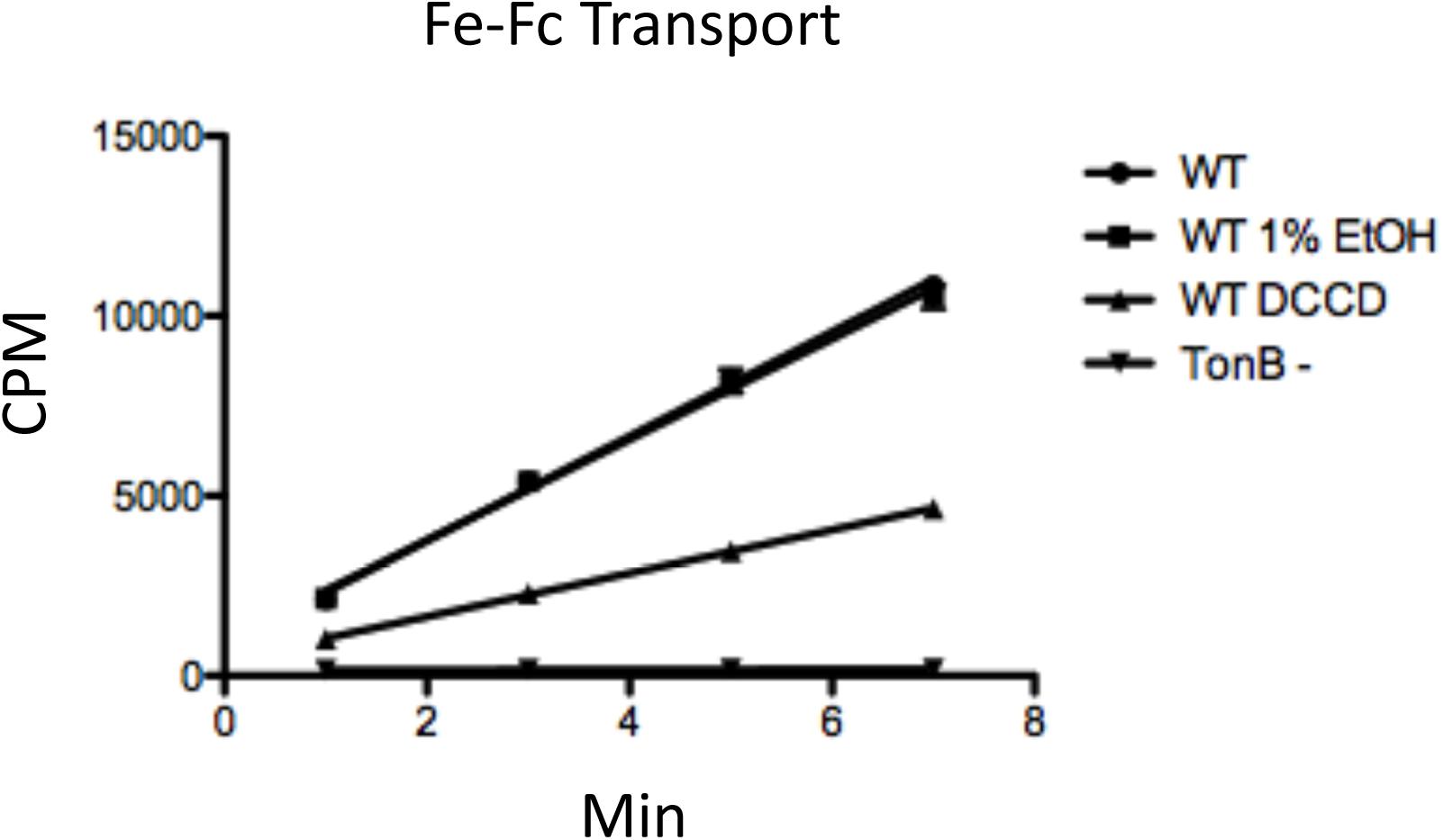
DCCD inhibits TonB system function. Either 10mM DCCD in ethanol or the equivalent amount of ethanol was added to assay medium immediately prior to measuring the initial rate of Fe^55^-ferrichrome transport as described in Materials and Methods. WT stands for W3110. TonB-stands for strain KP1477 (W3110 Δ*tonB*, P14::*kan*).

While *in vivo* DCCD treatment had no effect on the migration of monomeric TonB or FepA in immunoblots, it did modify monomeric ExbB and ExbD (Fig. 14). About half of the ExbB formed an apparently smaller-mass monomer (ExbB_L_ for “lower”, red asterisk). To test if ExbB_L_ was a proteolytic fragment, cells expressing ExbB variants with a carboxy terminal T7 epitope tag (ExbB-T7) or an amino terminal His_6_ epitope tag (His-ExbB) were treated with DCCD (Fig. 15). The ability to detect both epitope tags indicated that ExbB_L_ represented full length ExbB rather than a proteolytic fragment. The decrease in apparent mass of ExbB_L_ was thus likely due to a DCCD-mediated intramolecular cross-link similar to those caused by ExbB (pBpa) cross-linking shown in Fig. 8, lanes 8 and 10. Intramolecular cross-linking of ExbB could inhibit the TonB-dependent energy transduction cycle by preventing required conformational changes (65).

**Figure 14.**
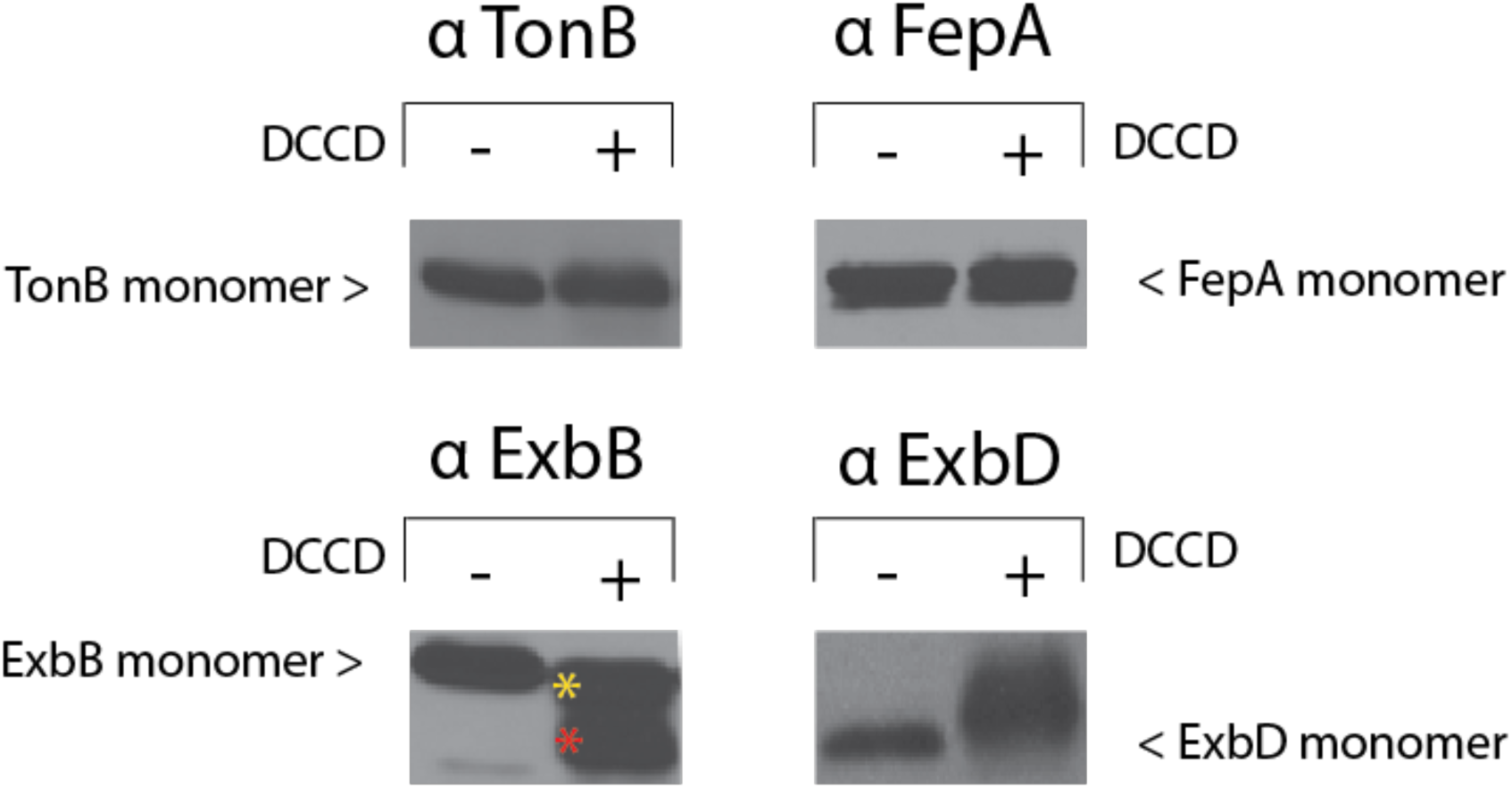
DCCD treatment causes aberrant migration of ExbB and ExbD in SDS-polyacrylamide gels. W3110 cultures were grown to mid-exponential phase at which point cells were harvested and resuspended in 1ml 100mM sodium phosphate buffer at pH 6.8. Either 10 µl of 100 mg/ml DCCD in ethanol (+) or 10 µl ethanol alone (-) was added to cells followed by incubation for 15 min at room temperature. Monomeric *p*-formaldehyde (final concentration = 1.0%) was subsequently added for a further 15 min incubation. Equal numbers of cells were processed for electrophoresis on 13% SDS-polyacrylamide gels and immunoblots developed with anti-TonB (top left), anti-FepA (top right), anti-ExbB (bottom left), or anti-ExbD (bottom right) antibodies (6, 92). The yellow asterisk identifies ExbB, and red asterisk identifies ExbB_L_ (lower). DCCD had no effect on previously characterized formaldehyde crosslinking profiles at higher masses [(19, 57,61); data not shown].

**Figure 15:**
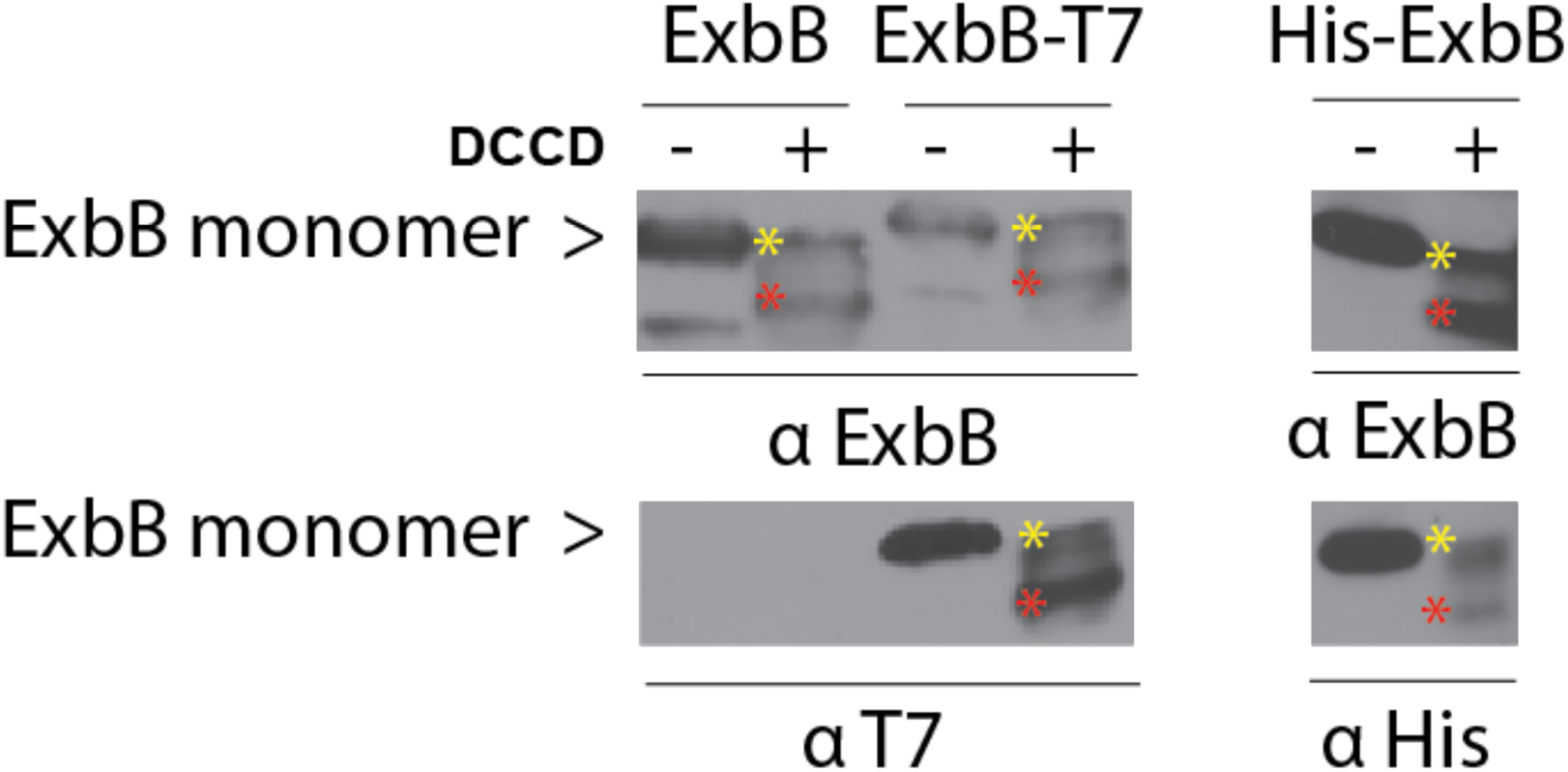
ExbBL (lower) is full length ExbB protein. In strain RA1017, cultures expressing plasmid-encoded ExbB (pKP660), ExbB with a carboxy terminal T7 epitope tag (pKP735) and ExbB with an amino terminal His6 epitope tag (pKP560) were either DCCD-treated (+) or ethanol-treated (-), then formaldehyde-cross-linked and immunoblotted as in Fig. 14, except that a final concentration of 1 mM DCCD was used. Immunoblots were developed with anti-ExbB (upper panels) with the identities of the samples indicated across the top of the panel. The same samples were also developed with anti-T7 tag (bottom left panel) or anti-His tag antibodies (bottom right panel). The position of monomeric ExbB is indicated on the left. For ExbB from DCCD-treated samples, the yellow asterisk identifies ExbB monomer and red asterisk identifies ExbB_L_ (lower). The two panels on the left show immunoblots from identical, divided samples developed with different antibodies; the two panels on the right show immunoblots from identical, divided samples developed with different antibodies.

DCCD treatment caused a majority of ExbD to form a smear of higher mass monomers *in vivo,* possibly representing modification of an array of conformations as it moved through an energy transduction cycle (Figs. 2, 14, and 16, lanes 1 and 2). To test this idea, we examined the effect of DCCD on a form of ExbD that is stalled at Stage II in the energy transduction cycle, ExbD (D25N) [Fig. 2; (45, 47)]. Treatment of ExbD (D25N) with DCCD caused the higher mass smear to collapse into a single higher mass form as well as an increased amount of monomer, suggesting that, at Stage II, only a single DCCD-modifiable conformation was available (Fig. 16, compare lanes 2 and 4). These results suggested that the various DCCD modifications could have stalled wild-type ExbD at several different Stages in the energy transduction cycle, resulting in its inactivation. They also identified a new assay with which to visualize ExbD conformational changes *in vivo*.

**Figure 16:**
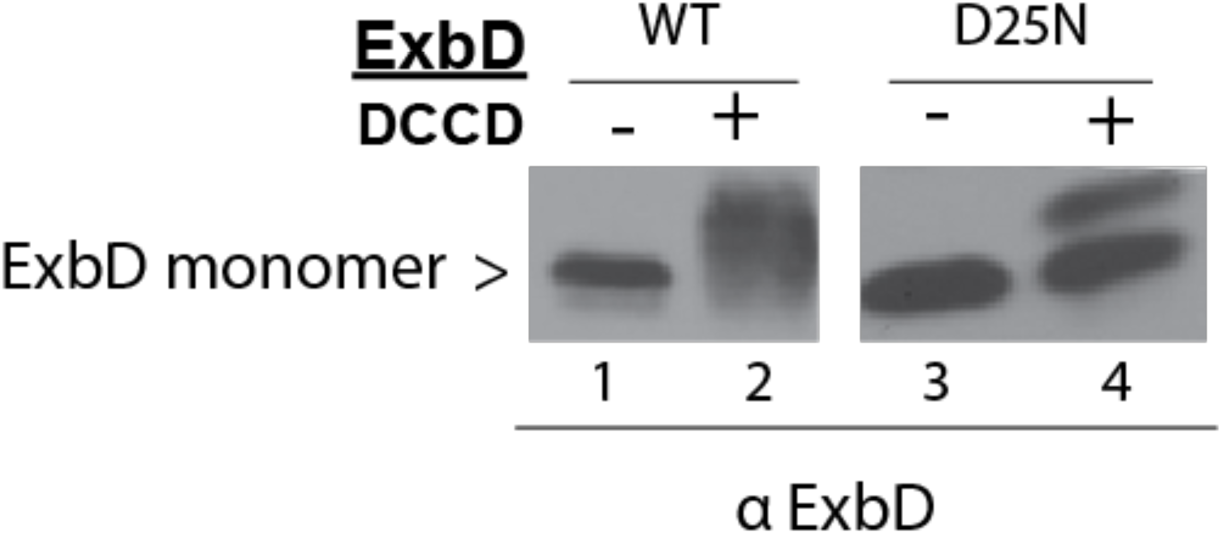
ExbD (D25N) resolves the smear of DCCD-treated ExbD into monomer and a single modified form. In strain RA1017, cultures expressing plasmid-encoded ExbD (pKP660), and ExbD D25N (pKP962) were either DCCD-treated (+) or ethanol-treated (-), then formaldehyde-cross-linked and immunoblotted as in Fig. 14, except that a final concentration of 1 mM DCCD was used. Equivalent numbers of cells were loaded on a single 13% SDS-polyacrylamide gel and immunoblotted. Immunoblots were developed with anti-ExbD antibodies (6). Lanes 1, 2, 3, and 4 are from the same immunoblot with intervening lanes removed. The position of monomeric ExbD is indicated on the left.

## DISCUSSION

### The cytoplasmic domains of ExbB bind unknown proteins

The bulk of the scaffolding protein ExbB occupies the cytoplasm with an ∼90 residue cytoplasmic loop and an ∼50 residue carboxy terminus [(31), Fig. 1B]. Given its membrane topology, the cytoplasmic residues of ExbB are a logical site of interaction with both known and unknown proteins. Here we showed that three out of six residues substituted with pBpa within the ExbB cytoplasmic loop formed photo-cross-linked contacts with an unknown protein of ∼34 kDa *in vivo*. Given that four of nine 10-Ala substitutions exclusive of those pBpa substitutions were inactive, there may be additional contacts with the ∼34 kDa protein to be discovered. In addition, photo-cross-linking through R222 (pBpa) in the ExbB carboxy terminal tail formed contacts with unknown proteins of ∼29 kDa and ∼24 kDa.

Our working model requires the existence of cytoplasmic proteins to assemble and move the proposed complexes of TonB_2_-ExbB_4_ and ExbD_2_-ExbB_4_ together and apart through the Stages that have been defined by *in vivo* experiments (Fig. 2). While we do not know the identities of these proteins, we now know for the first time that they exist. In that sense, the unknown cytoplasmic proteins might function like those that power adventurous motility, the paralogous AglQRS proteins in *Myxococcus*. (40, 76). Because the genes for unknown proteins have not yet surfaced in *tonB* system selections, the unknown proteins might be redundant or, given a possible link to growth regulation (28), essential. Characterization of the implicated proteins is likely to yield new insights into TonB-dependent energy transduction.

### ExbB cytoplasmic loop and tail domain variants mediate immediate growth arrest

The phenomenon of immediate growth arrest induced by overexpression of Δ10 ExbB variants, but not by wild-type ExbB, is striking (28). Because the plasmid-encoded Δ10s in the cytoplasmic loop of ExbB are proteolytically unstable, it is necessary to use high levels of inducer to achieve chromosomally encoded levels and characterize their properties.

Our previous study of the Δ10 deletions in the ExbB cytoplasmic loop shows a complete correlation of immediate growth arrest with proteolytic instability (28). While we postulated that immediate growth arrest is due to changed interactions with growth regulatory proteins, that correlation also raised the possibility that a fragment of ExbB, common to all, might inhibit a central growth pathway. However, even though four of the corresponding cytoplasmic loop 10-Ala substitutions from this study also required maximal levels of inducer to reach chromosomal levels—and were thus judged to be proteolytically highly unstable—none caused immediate growth arrest. Therefore, structural distortions caused by changed interactions of Δ10s with one or more growth-regulatory proteins were more likely the cause of the immediate growth arrest (28).

There may have been a different position-specific mechanism of immediate growth arrest mediated by the Δ10 and 10Ala substitutions in the cytoplasmic carboxy terminal tail. All three Δ10 deletions closest to transmembrane domain 3 caused immediate growth arrest, but as 10-Ala substitutions only the single ExbB Q215-L224 Ala did (Fig. 4 B, Table 3). Either of the two Δ10s closest to transmembrane domain 3 relocated ExbB Q215-L224 out of its correct position. Consistent with that, the two Δ10 s distal to ExbB Q215-L224, preserved its correct position relative to transmembrane domain 3 and did not exhibit immediate growth arrest. We hypothesize that the ExbB Q215-L224 Ala substitution triggered immediate growth arrest because, out of the entire ∼50 residue carboxy terminus, it alone lacked the correct residue, R222, to bind to growth regulatory proteins— likely the ∼24 kDa and ∼29 kDa unknown proteins.

Because each of the three cytoplasmic carboxy terminal tail Δ10s and the single 10-Ala that caused immediate growth arrest was also proteolytically unstable, the possibility that a proteolytic fragment of ExbB caused immediate growth arrest could not be ruled out for this admittedly smaller sample size. In either case, taken together, the results suggested that both the cytoplasmic domains of ExbB were involved in interactions with unknown proteins required for cell growth.

### *In vivo* and *in vitro* views of ExbB, ExbD, and TonB

There are similarities and differences between solved structures of ExbB_5_/ExbD_2_ and *in vivo* data (42, 45, 57). In the solved structures, ExbB is a pentamer with homodimerized transmembrane domains of ExbD occupying the entirety of its lumen (77, 78). Like the solved structures, the *in vivo* studies also show that ExbD homodimerizes through its transmembrane domain (45); unlike the solved structures, ExbB forms a dimer of dimers, a tetramer, *in vivo* (38). The *in vivo* studies also show that ExbB formaldehyde cross-links to both TonB and ExbD *in vivo*, likely through their transmembrane domains [(19, 57); Fig. 1]. However, TonB is absent from the solved ExbB_5_/ExbD_2_ structure. Because TonB and ExbD periplasmic domains clearly interact *in vivo* (51), monomeric TonB is hypothesized to bind the external surface of the solved ExbB pentamer via its single transmembrane domain (79–81).

The results we report here provided new data about *how* the two views of the TonB system differ. First, they are different because we found that the TonB transmembrane domain formed homodimers *in vivo*, which provided support for our model in Figure 2 that posits a homodimeric form of TonB stabilized by an ExbB tetramer. Thus, if nothing else, the structural models might be updated such that there are two homodimerized TonB transmembrane domains postulated to exist on the external surface of an ExbB pentamer, making the ratio in the structural models ExbB:ExbD:TonB 5:2:2. That ratio is significantly different than the ratio of 7:2:1 observed *in vivo*, with early studies of the paralogous Tol system finding similar *in vivo* ratios for TolQ:TolR:TolA (6, 82, 83).

Second, our results suggest that the TonB transmembrane domain homodimer may not be located on the external surface of the ExbB pentamer. *In vivo* photo-cross-linking showed here that the TonB transmembrane domain came within 3 Å of ExbD--almost certainly its transmembrane domain given the membrane topologies of the two proteins—at some point in the energy transduction cycle (Fig. 1). This result extended the reciprocal data that the ExbD transmembrane domain photo-cross-links to TonB at an unknown site (57) and supported Stage III of the model where data show that Asp25 in the ExbD transmembrane domain is required for correct interaction between periplasmic domains of ExbD and TonB (Fig. 2). In the solved structures, the lumen of the ExbB pentamer can accommodate only two transmembrane domains (77, 78).

This study has also provided new insight into *why* the two views of ExbB might differ. *In vivo* ExbB is a dimer of homodimers—a tetramer instead of the pentamer seen in solved structures (38). Here we found that unknown proteins, which the solved structures do not include, bound to cytoplasmic domains of ExbB *in vivo*. The presence of the cytoplasmic membrane and PMF could also influence protein-protein interactions that occur during TonB-dependent energy transduction *in vivo*. It is known that TonB interactions with the TBDT BtuB differ in whole cells compared to *in vitro* reconstitutions (84).

The dynamic, and possibly rapid, conformational changes through which the components of the TonB system appear to function could make it challenging to capture the full range of known interactions in structural studies (Fig. 2). The solved ExbB_5_/ExbD_2_ structures most closely resemble the ExbB_4_-ExbD_2_ complex from Stage I (Fig. 2). The *in vitro* and *in vivo* models of ExbB/D structure could be reconciled if the newly detected unknown cytoplasmic ExbB-binding protein(s) were required to configure ExbB as a tetramer (rather than pentamer), and if structural studies had not yet captured other configurations suggested in Fig. 2, such as ExbB_4_-TonB_2_, also from Stage I.

### Visualization of a PMF-dependent ExbD conformational change *in vivo*

The results in this paper eliminated the possibility that ATP synthase played a role in the TonB system. Instead, they suggested that the inhibitory effect of the ATP synthase inhibitor DCCD on TonB system activity was due to modification of ExbD and/or intramolecular cross-linking of ExbB *in vivo*.

The serendipitous discovery of ExbD modification by DCCD demonstrated *in vivo* structural changes in ExbD for the first time. In our current model the periplasmic domain of ExbD is hypothesized to undergo PMF-dependent conformational changes *in vivo* based on changes in sensitivity to exogenously added proteases and on identification of both ExbD homodimers and ExbD-TonB heterodimers, implying conformational differences; however, the Stages in ExbD behavior have not been detected outright [Fig. 2; (46, 47, 51, 52)]. By comparing DCCD-mediated modification of wild-type ExbD at a variety of sites with that of the ExbD (D25N) variant stalled at Stage II and modified at a single site, we were able to visualize differences in ExbD conformation proposed by the model (Fig. 2).

## MATERIALS AND METHODS

### Strains and plasmids

*Escherichia coli* K12 strains and plasmids used in this work are listed in Table 4. RA1016 was constructed from W3110 by the same method used for RA1017 (68), the λ-red recombinase approach of Datsenko and Wanner (85). The TolQ-phenotype was confirmed as insensitive to A-group colicins, sensitive to B-group colicins, and sensitive to detergent. RA1016 was P1*vir*-transduced with the *tonB*::*kan* mutation from the Keio collection (86) to create KP1564. The kanamycin cassette was removed from KP1564 using plasmid-encoded flippase of pCP20 (85) to create KP1565. The Δ*exbB/D*::*kan* mutation was P1*vir*-transduced from RA1003 (68) into KP1565 to create KP1566. KP1441 was created by P1vir transduction of the Δ*uncB-C* deletion using the linked *ilv*::Tn*10* in CK1801(87) into GM1 (88) and screening for inability to grow on succinate. Lack of ATPase subunits was confirmed by immunoblot (data not shown).

In-frame Δ10 deletions and 10-Ala insertions in *exbB* were created in plasmid pKP660 using extra-long PCR as described (28). Amber mutations in *exbB* were created in plasmid pKP1657 using extra-long PCR. Amber mutations in *tonB* were created in plasmid pKP315 using extra-long PCR. The *exbB* amino terminal His6-tag in pKP560, the *exbB* carboxy terminal T7 epitope tag in pKP660 and the D25N mutation in *exbD* of pKP962 were all created using PCR. The DNA sequences of the entire *exbB/D* operon or *tonB* gene for each plasmid were verified at the Penn State Genomics Core Facility (University Park, PA) and the existence of unintended base changes was ruled out.

### Media and culture conditions

Luria-Bertani (LB), tryptone (T), and M9 minimal salts media were prepared as described previously (89, 90). Except for plasmid-less strains, all liquid cultures, agar plates, and T-top agar were supplemented with 100 µg of ampicillin/ml with plasmid-specific amounts of arabinose added as inducer to approximate chromosomal levels seen in W3110. The exception was for pKP1657-based plasmids where propionate was used as the inducer to achieve chromosomally encoded levels. M9 salts were supplemented with 0.4% glycerol, 0.2% casamino acids, 40 μg/ml tryptophan, 4 μg/ml thiamine, 1 mM MgSO_4_, 0.5 mM CaCl_2_, and 37 μM FeCl_3_. Cultures were grown at 37°C with continuous aeration.

In vivo *photo-cross-linking*. Plasmid pEVOL encodes an orthogonal tRNA synthetase and a corresponding orthogonal tRNA_CUA_ that recognizes amber (UAG) codons (58). Plasmids with amber substitutions in *exbB* or *tonB* were engineered, co-expressed along with pEVOL in various strain backgrounds, and processed as previously described (57).

### Initial rates of [^55^Fe]-ferrichrome transport

The initial rates of iron transport for ExbB Δ10 and 10 Ala variants in Tables 1 and 2 were determined as described previously (28), where mid-exponential cultures were induced with the levels of arabinose required to achieve chromosomally encoded levels of expression, incubated for 60 min and equal numbers of cells subsequently assayed for initial rates of iron transport. The equivalence of cell numbers assayed was confirmed by staining immunoblots for total protein with Coomassie stain. For the data in Fig. 13, mid-exponential-phase subcultures were pelleted and suspended in iron transport assay media, and the initial rates of [^55^Fe]-ferrichrome transport were determined as described previously following addition of 1% ethanol or 10mM DCCD (N, N_1_-dicyclohexylcarbodiimide) in ethanol (91).

### Immunoblot analysis

Evaluation of *in vivo* protein levels was carried out by immunoblotting to PVDF (polyvinylidene fluoride) membranes and development with anti-ExbB, -TonB, and - ExbD antibodies, as described in (6, 91, 92). Following development of the immunoblot by chemiluminescent detection with secondary antibodies, the PVDF membranes were stained with Coomassie blue to control for equal loading of the samples. Anti-His-specific monoclonal antibodies were from EMD Millipore (prod # 70796). Anti-T7 tag-specific monoclonal antibodies were from Novagen.

## ACKNOWLEDGEMENTS

We thank James Fisher, Erica Ward, Glenn Hwang, and Noriko Mikeasky for *excellent* technical assistance; Gail Deckert for construction of pKP560; Mary Huber for construction of pKP962; Corinna Moro for construction of Δ10 mutations in the ExbB carboxy terminal tail; Kristin Baker for construction of KP1566; Jacob Mowery and Avinash Bakshi for construction of ExbB amber mutants; Aleena White for construction of TonB amber mutants; and Fanny Kippelen for sequence comparisons of the ExbB carboxy terminal tail. We thank Penelope Higgs for providing anti-MreB antibodies and Karlheinz Altendorf for providing anti-ATPase subunit antibodies. We thank Ray Larsen for critical reading of the manuscript. Support from NIAID R21 AI113622, NIGMS R01 GM112710, NIGMS R01 GM42146 and Violet S. Postle is gratefully acknowledged.

